# Anti-cancer property of *Lenzites betulina* (L) Fr. on cervical cancer cell lines and its anti-tumor effect on HeLa-implanted mice

**DOI:** 10.1101/540567

**Authors:** Tapojyoti Sanyal, Swapan Kumar Ghosh

**Affiliations:** Molecular Mycopathology Lab, Cancer Research Unit, PG Dept. of Botany, Ramakrishna Mission Vivekananda Centenary College (Autonomous), Rahara, Kolkata 700118, India

**Keywords:** *Lenzites betulina*, ethanolic extract, HeLa, SiHa, CaSki cell, cervical cancer, tumor, cytotoxicity, apoptosis, migration, colonization, autophagy, metastasis, tumor, mice

## Abstract

In global scenario cervical cancer is increasing. New drugs from natural compounds are in search. Mushrooms are now recognized as miniature pharmaceutical factories producing hundreds of novel constituents. We have taken ethanolic extract *Lenzities betulina* (LBE) wild mushroom for evaluation of its as anti-cancer property against cervical cancer cell lines e.g. HeLa, CaSki and SiHa and anti tumor activity against HeLa implanted tumor on mice. The extraction was done by dip and stirring method in 90% ethanol for 72 h. For evaluation of anti-cervical cancer, several assays were performed such as MTT assay, cell morphology by phase contrast microscope and F-action polymerization by Laser scanning confocal microscope and nuclear morphology DAPI staining under inverted fluorescence microscope, MMP, ROS, cell cycle, autophagy and stem cell population by flow cytometry and DNA laddering were done. Western blotting was done for protein expression. To evaluate anti-metastatic activity, anti-cologenic assay and wound healing assay were adopted. For chemo-analysis of the LBE, GC-MS was done. The results from Cytotoxicity assay showed that at highest dose of LBE (1000 µg/ml) after 24 h, percentage of cell inhibitions were 85.13 %, 77.13 % and 47.70 % against HeLa, CaSki and SiHa respectively and the calculated IC_50_ values were 492.52 ± 2.6 µg/ml, 612.22 ± 4.2 µg/ml, and 1210.30 ± 6.4 µg/ml respectively. Depending upon the cytotoxicity screening, HeLa cell line was considered for the further studies. Cell morphology study exhibited that LBE treated HeLa cells became round from normal spindle shape. DAPI staining showed that LBE treated nucleus became condensed and fragmented. DNA fragmentation at 230 and 300 base pair zone from agarose gel assay was observed. LBE induced ROS generation and reduced MMP. It up regulated the expression of apoptotic genes and p53 while down regulated Bcl2, pro-caspase 3 and pro caspase-9 gene. Cell cycle was arrested at G2/M checkpoint. Autophagic induction was exhibited by vacuole formation in treated cells. CSC population of treated cells was reduced and F-actin polymerization was observed in treated cells. In addition, LBE suppressed metastatic nature by inhibition of cell migration and colonization. The inhibition of growth of the tumors in HeLa cell-implanted mice showed that treatment with 50 mg LBE/kg of body weight of mice led to a marked reduction in the volume (93.22 ± 9.2 %) and weight (90.42 ±9.55 %) of the tumors. The GC-MS profile of LBE shows that out of 69 compounds, 9, 12-Octadecadienoic acid (Z, Z) and Ergosta-5, 8, 22-trien-3-ol, (3.beta22E) are in a significantly higher proportion with the percentage peak area 22.13 and 19.72 respectively. Library search for bioactivity showed that these compounds are anti-cancerous and interestingly 4’-Hydroxy-6-methoxyaurone binding with P-glycoprotein inhibits the cancer cells to become drug resistant. In conclusion, LBE is very prominent anti-cervical cancer having a lot of anti-cancerous compounds which are probably acting synergistically. This report of anti-cervical cancer property of *L. betulina* is probably first time in oncology. Its therapeutic use in human model is urgent for new drug development.

## INTRODUCTION

Mushrooms or macrofungi offer high sources of new isolable bioactive compounds with diversified chemical structures, which are considered potent sources for drug discovery. In the stone era, human went for hunting and the same time collect food including mushrooms for organoleptic properties such as flavor and texture ^1-3^and then realized that some of mushrooms have properties for curing certain ailments. Local consumption of wild mushrooms are also increasing because of their medicinal properties^4, 5^due to the presence of secondary metabolites having pharmaceutical importance.^3, 4^ There are more than 14,000 mushrooms out of 5.1 million estimated fungi, among which near about 700 exhibit medicinal properties^6^. *Lenzitis betulina* (L) Fr. belongs to the family *Polyporaceae,* order *Polyporales* and phylum to *Basidiomycota,* and grows on logs and fallen woods. Various species of *Lenzites* are well distributed around the world. *Lenzitis* spp have no value as food since it has an extremely tough exterior^7^. There have been different studies on the various medicinal uses of *Lenzites* spp. antimicrobials^8, 9^, anti-viral^10^, and immunosuppresser^11^. Phytochemical analysis of compounds presents in *Lenzites* spp. using three different solvents (ethanol, water and petroleum ether) showed that phenolic and steroids were present in the ethanol extract; flavonoids, tannins and steroids were present in the petroleum ether and only saponins were present in the aqueous extract^9^. Since many of the compounds have been shown to act synergistically, it is worth testing the anti-proliferative effects of the whole mushroom extract rather than its individual components. This principle (synergy) is compatible with similar natural biological products like the essential oils, which are more effective when used as whole products, while quenching or nullifying potential unwanted side-effects by the presence of individual components^12^. Among mushroom extracts, an ethanol extract probably finds the most extensive application^13^. Cancer **i**s a major public health problem and one of the leading causes of death in the world today^14^. Cervical cancer is the second most common cancer among women worldwide^15, 16^. One third of newly diagnosed cancers among women in the India are cervical cancers and that is for the infection with some types of HPV (Human Papilloma Virus). Every year in India, 122,844 women are diagnosed with cervical cancer and 67,477 die from the disease^17 18^. The current anti-cancer drugs i.e. chemo-drugs available in market are not target specific and pose several side-effects and complications in clinical management of various forms of cancer^19^, which highlights the urgent need for novel effective and nontoxic natural compounds^20,21^. Epidemiological studies showed that among the Chinese women who regularly intake powder of *Agaricus bisporus* with tea have less breast and gastrointestinal (GI) cancers^22^. Consumption of mushroom has inverse effect in breast cancers occurrence among premenopausal women^23^. Lucas *et al.*^24^got the first credit to show that mushroom extract has antitumor properties using extracts of *Boletus edulis* and other basidiomycetes against Sarcoma-180 and also on HeLa cell lines presenting significant growth inhibition on those types of cancer. Ikekawa *et al*^25^ published one of the first scientific reports on antitumor activity of extracts of mushrooms against implanted Sarcoma 180 in animals. Soon after, three major anticancer drugs, Krestin from cultured mycelium of *Trametes* (*Coriolus versicolor*), Lentinan from fruiting bodies of *Lentinus edodus* and Schizophyllan from *Schizophyllum commune*, were developed ^26-28^. We came to know from the review work of Dai Yu-C *et al* ^29^, that 200 to 331 of mushroom species have anticancer activity. Near about 650 species of higher basidiomycota have been found anti-cancer activity^26^. Although *Lenzites betulina* is worldwide distributed, there has been minimal or no extensive research work on it for cancer remedy. Therefore, the main objectives of this work are to screen cytotoxicity effect of ethanolic extract of *Lenzites betulina* (LBE) aganist HeLa,CaSki and SiHa cell lines of cervical cancer, to evaluate anti proliferative, apoptotic, autophagic activity, effect on cancer stem cell and anti-metastatic (anti migration /anti-invasion and anti-colonization) activity of LBE on HeLa cell line, to evaluate antitumor effects of LBE in HeLa-implanted mice and to find out chemo profile by Gas chromatography and mass spectrometry (GC-MS).

## MATERIALS AND METHODS

### Materials

The fruit bodies of *Lenzites betulina* (L) Fr were collected from dead logs from Baruipure, South 24 Parganas, West Bengal of India and carried to laboratory by biodegradable carry bag and identification was done by morphological and anatomical analysis consulting with mushrooms keys, manuals and books^30-32^. One voucher specimen was kept in Fungal Herbarium of Department of Botany, Ramakrishna Mission VC College, and Rahara. Remaining were washed by running tap water, blotted by blotting papers and kept at 50^°^ C in oven for 48 h for drying.

### Preparation of ethanol extract (LBE)

The collected and identified fruit bodies of *L. betulina* (LB) after oven dried at 50^°^C temperature for 48 h were ground by mixture grinder (Beckman food mixer, India). This smashed biomass (100 g) was suspended in 500 ml of 90% ethanol and incubated for 72 h at 200 rpm and 37°C. The suspension was filtered on what man No. 4 paper to remove the biomass. This procedure was repeated thrice. The supernatant was concentrated and ethanol was subsequently removed from the extract using a rotary vacuum evaporator at 40°C, and the remaining solvent was removed with a freeze-drier. The resulting dried powder stored at 4°C. The stock solution of LBE(LBE) was prepared by dissolving in dimethylsulphoxide (0.2% DMSO) at a concentration 10 mg/ml and stored at 4°C.

### Cell culture

HeLa, CaSki and SiHa human cell line obtained from NCCS Pune were maintained in Dulbecco’s modified Eagle’s medium (DMEM) containing penicillin (50 U/ml), streptomycin (50 U/ml) and 10% fetal bovine serum (FBS). The cultures were maintained at 37°C in 5 % CO_2_ and 95% humidity in CO_2_ incubator (Thermo scientific, USA).

### Chemicals

3-(4,5-dimethyl thiazol-2-yl)-5-diphenyltetrazolium bromide (MTT), Fetal Bovine Serum(FBS), Phosphate Buffered Saline (PBS), Dulbecco’s Modified Eagle’s Medium (DMEM), and Trypsin were obtained from Himedia Laboratory. EDTA, glucose, and antibiotics from Himedia Laboratories, Dimethyl Sulfoxide(DMSO) from Cell Clone and Propanol from Thermo Fisher Scientific. JC1 (Thermo Fisher Scientific), RIPA (Radioimmunoprecipitation assay buffer), H_2_DCFDA (Thermo Fisher Scientific) L-ascorbic acid (SRL), DPPH (SRL) H_2_O_2_ (Merck). Annexin-V kit from Santacruze Biotechnology,USA. Anti-mouse antibodies against p53, Beta actin, Bcl-2, pro-caspase-3, pro-caspase-9, PARP, were procured from Santa Cruz USA Nitrocellulose membrane and filter papers were obtained from Bio-Rad. Dapi(Thermo Fisher Scientific) Ethanol were HPLC grade and were purchased from Merck. Alexa Fluor 488 Phalloidin and Hoechst 33342 (Thermo Fisher Scientific). The others chemicals and materials were purchased from local firms (India).

### Cell lines, culture medium and subculture of cell line

Human Cervical Cell lines (HeLa, CaSki and SiHa) were grown and maintained in DMEM (Dulbecco’s Modified Eagle’s) Medium supplemented with L Glutamine, 10% FBS, sodium bicarbonate in T25 cell culture flasks to the 80-90% confluence. The media also supplemented with 1000 μg/ml streptomycin (Himedia) and 1000 IU/ml penicillin (Himedia) at 37°C in a humidified atmosphere of 5% CO_2_. Maintenance cultures were passaged weekly, and the culture medium was changed twice a week^33^.

### Cytotoxicity / cell proliferation assay

MTT colorimetric assay method was employed to evaluate cell viability in this cytotoxic assay. Cells of HeLa, SiHa and CaSki were seeded in a 96 well plate (1×10^4^ cells/well in 100 μl of medium) separately and treated for 24, 48 and 72 h with 100, 250, 500, 750 and 1000 µg/ml LBE. After incubation the media was discarded and 100 μl/well MTT (Himedia Laboratory) reagents were added into each well and the cells incubated for another 2-4 h until purple precipitates were clearly visible under a microscope. Flowingly, the medium together with MTT (100 µl) were aspirated off the wells, DMSO (100 µL) was added and the plates shaken for 5 min. The absorbance for each well was measured at 540 nm in a micro-titre plate reader (Bio-Rad i-mark). The IC_50_ values of LBE were calculated

The concentration which led to a 50% killing (IC_50_) was calculated by plotting a dose response graph of the cytotoxicity values obtained using the formula given below:

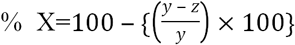, Where x= % of cell Cytotoxicity, y= Control, Z= test. Data points represent the mean ± SD in one experiment repeated at least thrice.

### Morphological examination of HeLa cancer cells

HeLa cancer cells were grown in 6 well-culture plate and treated with LBE (0, 500 μg /ml and1000 μg /ml). Cells were examined under phase contrast inverted microscope (Olympus,Japan) and photographs were taken using a digital camera (Q Imaging) for analysis.

### 4, 6-Diamidino-2-phenylindole (DAPI) nuclear staining

HeLa cells were cultured in cover slip in 6 well plates and treated with 500 and 1000 µg/ml of LBE and incubated at 37°C for 24 h. After incubation cells were washed with PBS in three times and the cells were fixed in 3.7% formaldehyde for 10 min then the cells were rinsed into PBS+ for three times, 0.2% Triton X-100 was added and incubated for 5 min. After incubation the cells were washed with PBS and DAPI was added, and incubated for 5 min, the solution was discarded and again the cells were washed with PBS in three times. After washing cells were observed under the fluorescence microscope (Olympus inverted fluorescence microscope CKX 53)

### Effect of LBE on the mitochondrial membrane potential (MMP) of HeLa cells

Mitochondrial outer membrane permeability was measured by flow cytometry using the MitoProbe(tm) JC-1 Assay Kit, Thermo Fisher Scientific. HeLa cells were cultured in 6 well plates and treated with 250 and 500 µg/ml of LBE and incubated at 37°C for 6 h and 12 h. After incubation cells were washed twice with PBS and incubated with 10 µl of 200 µM JC1 at 37°C, 5% CO_2_ for 30 min, cells were washed once by adding 2 ml of warmed phosphate-buffered saline (PBS) to each tube of cells. The cells were pelleted by centrifugation and re-suspended by gently flicking the tubes. 500 μl PBS was added to each tube and analyzed on a flow cytometer with 488 nm excitation using emission filters appropriate for Alexa Fluor 488 dye and R-phycoerythrin. A gate on the cells, excluding debris. Using the CCCP-treated sample, standard compensation intensity was determined by flow cytometry. Cells with reduced fluorescence were counted as having lost some of their mitochondrial membrane potentials.

### Detection of Intracellular ROS(Reactive oxygen species) by H_2_DCFDA

ROS were detected by the modified method of Deem TL, Cook-Mills JM.^6^ 1×10^5^ cells were cultured in 35 mm plates and treated with 500, 750 µg/ml of LBE and incubated at 37°C at 2 h. After incubation cells were trypsinized and suspended in 500µl PBS (1x), 5µl H_2_DCFDA (Thermo Fisher Scientific) was added and incubated for 30 min at 37°C.then analyze the sample in Flow Cytometer (FACS Calibur, BD Bioscience). Where Ascorbic acid acts as a positive control and H_2_O_2_ acts as a negative control. Also we observed ROS by inverted fluorescence microscopy.

### Cell cycle analysis

For the determination of cell cycle phase distribution of nuclear DNA. Cells were treated with 100, 250, 500 µg/ml LBE and incubate for 6 and 12 h. After incubation cells were fixed with 3% p-formaldehyde, permeabilized with 0.5% Triton X-100, and nuclear DNA was labeled with propidium iodide (PI, 125 mg/ml, Santacruze Biotechnology, USA) after RNase treatment. Cell cycle phase distribution of nuclear DNA was determined on FACS Calibur using Cell Quest software (Becton Dickinson). Histogram of DNA content (x-axis, PI fluorescence) versus counts (y-axis) has been displayed. Cell Quest statistics was employed to quantities the data at different phases of the cell cycle.

### Annexin V-FITC apoptosis assay

In general phosphatidyl-serine (PS) exists in internal face of plasma membrane but in early apoptosis it comes to external face of the plasma membrane, the high affinity PS binding protein Annexin V (conjugated to fluoresce in isothiocyanate FITC) which has high affinity to PS binding is used to identify the early event of HeLa cell killing caused by the LBE. On the other hand, simultaneously counterstaining propidium iodide (PI) is applied which allowed the differentiation of necrotic and apoptotic cells. An Annexin v-fluoresce in isothiocyanate (FITC)/ propidium iodide apoptosis detection kit (Santa cruze Biotechnology, USA), according to the manufacturer’s protocol, was applied to calculate cell apoptosis. In this study, percent of apoptotic and necrotic cells were determined after treatment with the LBE (250, 500, 750 and 1000 µg/ml) by flow cytometry using the AnnexinV-FITC apoptosis detection kit (Santacruze Biotechnology). Briefly, PI and Annexin-V were added to the cell line. The mixture was incubated for 15 min at 37°C, and then analyzed on FACS Verse. Electronic compensation of the instrument was done to exclude overlapping of the emission spectra. Total 10,000 events were acquired, the cells were properly gated and dual parameter dot plot of FL1-H (x-axis; FLUOS-fluorescence) versus FL2-H (y-axis; PI-fluorescence) was performed and shown in logarithmic fluorescence intensity.

### DNA fragmentation assay/ DNA Laddering assay

The DNA ladder assay was sensitive, cost-effective and most useful method for estimating apoptosis. 2×10^5^ cells were cultured into 6 well plates and treated with 500, 750 and 1000 µg/ml of LBE and incubate at 37°C for 48h. After treatment, DNA was isolated from HeLa cell line and visualized after the separation by gel electrophoresis (1.8% agarose gel containing Etbr in 40Mm Tris-acetate with electrophoresis at 75V at 4 h.)

### Autophagy assay by acridine orange

2×10^5^ cells were cultured in 6 well plates and treated with 250 and 500 µg/ml of LBE and incubated at 37°C at 24 h. After incubation cells were trypsinized and suspended in 500 µl PBS (1x), add 1µg/ml acridine orange and incubate for 30 min at 37°C. The sample was then detected using Flow cytometry (FACS Calibur, BD Bioscience) and data were analyzed

### F-actin staining

2×10^5^ cells (HeLa) were cultured in 6 well plates and incubated at 37°C at 24 h. After incubation cells were treated with LBE (750 µg/ml) and incubate at 37°C at 24 h. After incubation cells were washed with 1x PBS and fixed with 3.7% formaldehyde. After fixation 0.2% Triton X was added and incubated for 2 min and then Alexa flour 488 Phalloidin (1:500) was added for 1h in room temperature. Finally cells were stained by DAPI and the sample was then detected using confocal microscopy (Olympus, Japan) and data were analyzed.

### Hoechst efflux assay for cancer stem cell population study

2×10^5^cells (HeLa) were cultured in 6 well plates and treated with 250 and 500µg/ml of LBE and incubated at 37°C at 24 h. After incubation cells were trypsinized and suspended in 500 µl PBS (1x), verapamil 50 µg/ml was added and incubated for 30 min at 37°C. After verapamil treatment, cells were washed with PBS (1x) for 3 times, Hoechst (1µg/ml) was added and incubated for 30 min in 37°C. The sample was then detected using Flow cytometry (FACS Calibur, BD Bioscience) and data were analyzed.

### Western blot analysis

HeLa (2×10^5^) cells were treated with 100, 200, 500 and 750 µg/ml of LBE for 24 h. After treatment, cells were lysed with RIPA buffer (Abcam). The effect of treatment on the expression of certain cell cycle proteins such as p53, and on apoptotic proteins such as Bcl-2, pro-caspase-3, pro-caspase-9, and PRAP (Santacruze Biotechnology, USA) was determined. Proteins were detected by incubation with the corresponding primary antibodies, and antibodies followed by blotting with the HRP-conjugated secondary antibody. The blots were then detected using Luminol (Bio-Rad).

### Wound healing/ scratch assay

HeLa (50×10^3^) cells were seed in a 24 well plate and incubate at 37°C for 24 h. After 80-90% confluences, 200 μl micro tips used to press firmly against the top of the cell culture plate and a vertical wound down through the cell monolayer was swiftly made and then discarded the media carefully and each wall was washed with 1x PBS and finally treated with 500, 1000 µg/ml of LBE and a snap (zero h) through inverted fluorescence microscope (Olympus inverted fluorescence microscope, Japan) were taken and the culture plate was placed into CO_2_ incubator (37°C, 5% CO_2_). After 24 h incubation plates were removed and placed it in an inverted fluorescence microscope to take a snap (Olympus inverted fluorescence microscope, Japan) and sample was analyzed by Q Capture pro-7.

### Cologenic assay

HeLa (500) cells per well were seeded in a 6 well cell culture plate and incubated at 37 °C. Immediately after attachment the cells were treated with (250, 500,750, 1000 and 1250 µg/ml) LBE for 4 days of interval. Cells were treated with same concentration of extract then colonies were fixed with 3.7% formaldehyde and stained with (0.5% w/v) crystal violet. Viable colonies were counted using Olympus inverted fluorescence microscope (CKX 53). Colonies consisting of more than 50 cells were counted by using a colony counter and the results were reported as a percentage of colonies formed using the following equation: % Colonies formed = Colonies formed in treated sample /Colonies formed in untreated sample X100.

### Antitumor effects of LBE in HeLa cell-implanted mice

Male Swiss Albino mice (20-30 g) were used for the present investigation. The animals were kept in well ventilated cages (one group per cage) and provided with standard laboratory rodent diet *ad libitum* along with free access of water. Prior to commencement of experiments, all the animals were kept for four weeks under controlled temperature (25 ± 2 °C) and humidity (50-70%) with 12 h day/night cycle for acclimatization. All mice were housed under hygienic /pathogen-free air conditions in a room maintained at 24°C with 50% relative humidity and a 12 h/12 h light-dark cycle. All animal experiments were approved and performed according to the regulations of the Institutional Animal Ethics Committee, Government of India (approval no: 12/P/S/ IAEC/2018). The mice were divided into 6 groups:) control mice supplemented with pellet (control + vehicle; *n*=6);ii) controls supplemented with LBE (500 mg/kg of body weight) (control + LBE; *n*=6); iii) HeLa cell-implanted mice supplemented with pellet (HeLa + vehicle; *n*=6); iv) HeLa cell-implanted mice supplemented with LBE (10 mg/kg of body weight) (HeLa + LBE 10;*n*=6);v) HeLa cell-implanted mice supplemented with LBE (25 mg/kg of body weight) (HeLa + LBE 25, *n*=6); and vi) HeLa cell-implanted mice supplemented with LBE (50 mg/kg of body weight) (HeLa + LBE 50, *n*=6).

For tumor generation, a suspension of 2×10^6^ HeLa cells in 0.2 ml DMEM was subcutaneously injected on the dorsal surface of the right hind legs of the Swiss albino mice, where as the control groups were injected with DMEM. The tumors were measured with Vernier calipers every 3-4 days by using the formula *a*^2^ x *b*x 0.52 (where *a* is the shortest diameter and *b* is the longest diameter). When the tumor volume was measured to be 0.50-0.90 cm^3^, the mice were randomized. Subsequently, the mice were supplemented daily with vehicle or LBE at the doses of 10, 25, and 50 mg/kg of body weight for 28 days. After 28 days, all the mice in groups 3-6 were sacrificed, and the tumors were excised from their legs and then weighed. Visceral organs (i.e., the kidney, liver, spleen and lungs) were morphologically observed for possible side effects of LBE in the mice.

### Statistical analysis

All data were analyzed using Graph Pad Prism 5 software (Graph Pad Software, Inc., La Jolla, CA, USA), and one-way analysis of variance was performed to compare differences between the groups. All results are presented as the mean ± standard error of mean from three independent experiments performed in a parallel manner, unless otherwise indicated. All figures shown in the present study were obtained from at least three independent experiments. P<0.05 was considered to indicate a statistically significant.

### Gas chromatography-mass spectrometry (GC/MS) analysis

GC/MS analysis was done by Agilent Technologies, GC-6860N Network System with inert Mass Selective Detector and the search library used by C:\Database\NBS75K.L. Applying HP-5MS (19091S-602) Column the length and thickness of column was respectively30m and 0.25mm. The carrier gas was Helium gas (99.99%) used at a flow rate of 1ml/min and an injection volume of 2µl. Injector temperature was 280°C, 70ev ion source temperature 280°C. The oven temperature was 50°C isothermal for 2.0 min, with an increase of 10°C/min to 280°C, then10°C/min to 300°C, and ending with 10 min isothermal at 280°C.

## RESULTS

### Cytotoxicity / anti proliferative of LBE against HeLa cell line by MTT assay

It was investigated that whether LBE affected HeLa, CaSki and SiHa cell proliferation or not. The cell proliferation graph of the HeLa cell line (Fig1A, B, C) exhibited that untreated (negative control) cells proliferated at the expose time periods (24, 48 and 72 h). In contrast, cells were treated with LBE at all tested concentrations, showed reduced proliferation in concentration and time dependent manners in all cell lines (HeLa, CaSki and SiHa). Cells were treated with higher concentration (1000 µg/ml) of LBE proliferated least (14.87%, 22.87%, 52.30%) respectively. It indicated the percentage of cell inhibition were 85.13 %, 77.13 % and 47.70 % against HeLa, CaSki and SiHa (Fig 2A, B) at 24 h and 1000 µg/ml of LBE. In case of HeLa percentage of cell inhibition was increase as dose and time dependent manner. After calculating the data presented in graph 1, the IC_50_ value i.e. the concentration of LBE that resulted in a 50% reduction in absorbance compared with the control (negative) after 24 h of treatment, were 492.52 ± 2.6 µg/ml, 612.22 ± 4.2 µg/ml, and 1210.30 ± 6.4 µg/ml (HeLa, CaSki and SiHa) respectively.

**Figure 1.**
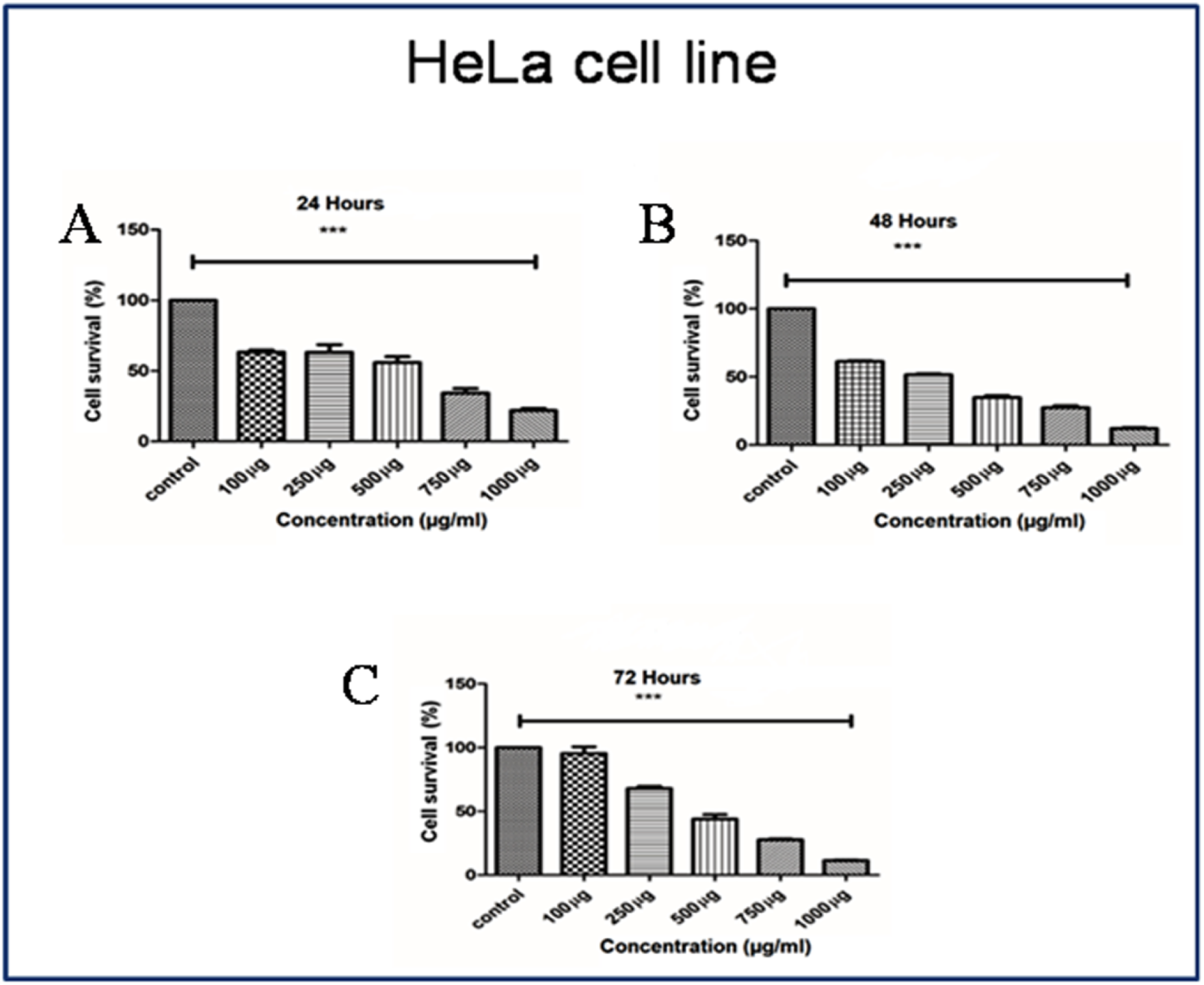
Cytotoxicity assay against HeLa cell line. (A) Cytotoxicity graph at 24h, (B). Cytotoxicity graph at 48 h, (C). Cytotoxicity graph at 72 h. Data are representative as a mean ±SEM of three independent experiments indicates *p<0.05, **p<0.01, ***p<0.001 as compared to control.

**Figure 2.**
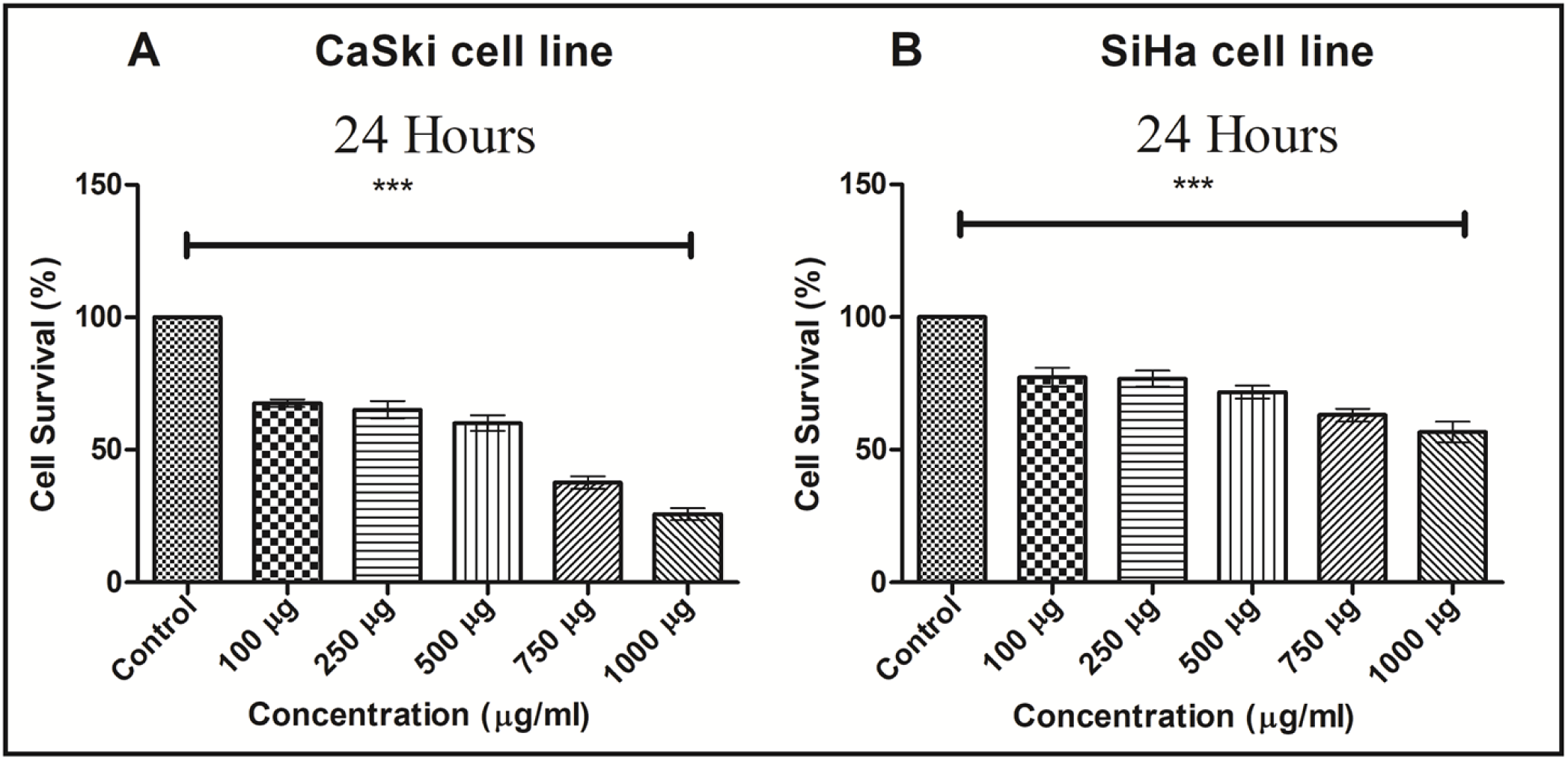
Cytotoxicity assay against SiHa and CaSki cell lines at 24 h (A) Cytotoxicity graph of CaSki cell line, (B) Cytotoxicity graph of SiHa cell line. Data are representative as a mean ±SEM of three independent experiments indicates *p<0.05, **p<0.01, ***p<0.001 as compared to control.

### Study of morphological changes of HeLa cells by LBE under Phase contrast microscopy and Fluorescence Microscopy

The cells, treated with various concentrations of LBE, and after 24 h cells, were examined under a phase contrast microscope for monitoring their morphology. Untreated cells (negative control) exhibited normal spindle shape and reached 90% confluency after 24 h culture. Cells, treated with 500 µg/ml and 1000 µg/ml of LBE were round and shrunken and cell membrane blebbing were observed in cells treated with 1000 µg /ml of LBE (Fig.3A, B and C). DAPI staining under fluorescence microscopy showed that increase of concentration of LBE, the intensity of fluorescence increased and nuclear shrinkage, fragmentation, chromosome condensation and apoptotic bodies, and the number of apoptotic cells increased in compare to the negative control (Fig D, E and F).

**Figure 3.**
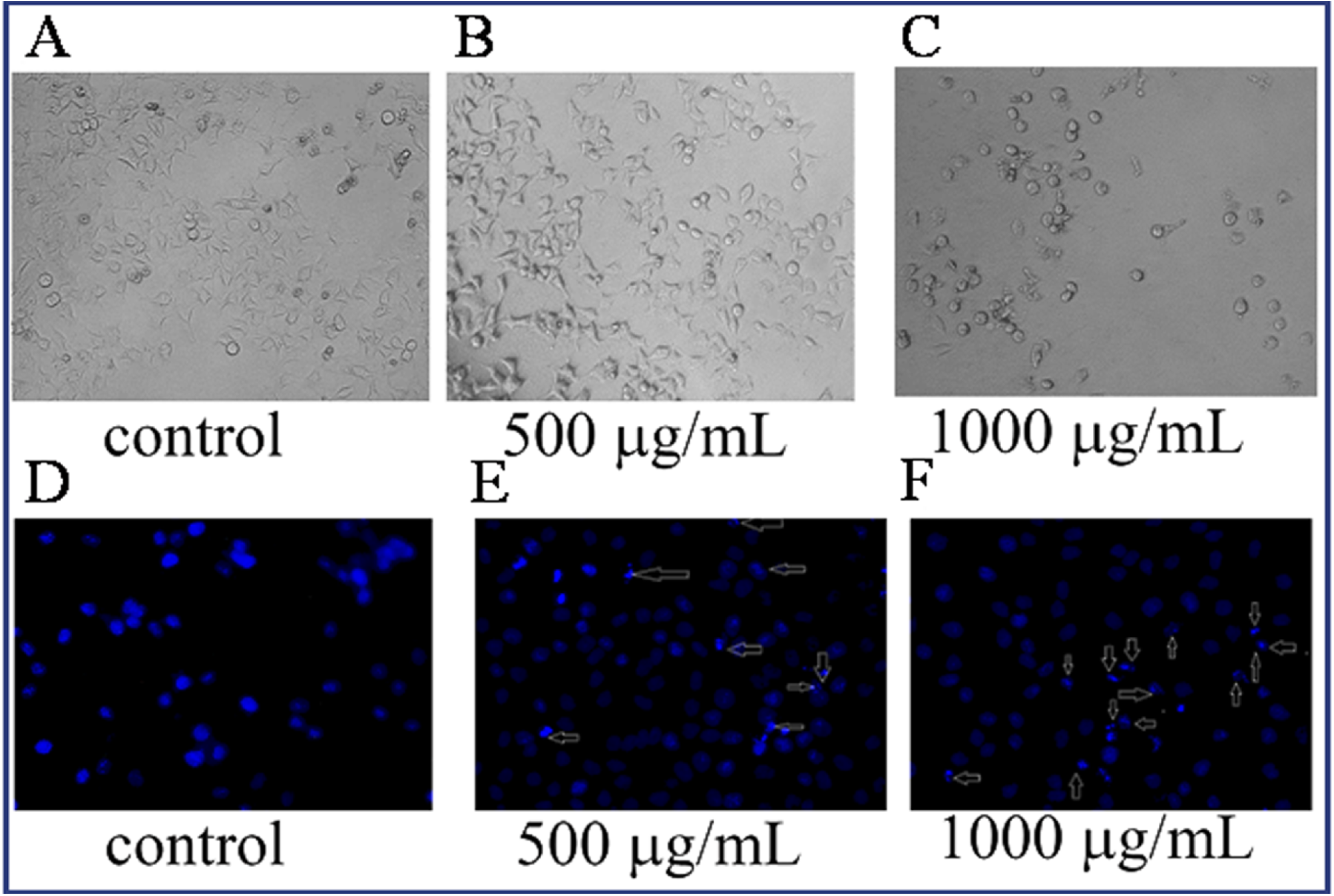
Cell morphology of untreated and treated cells. (A). Untreated (negative) HeLa cell, (B). LBE treated cells (500 µg/ml), (C). LBE treated cells (1000 µg/ml). Nuclear morphological study by DAPI staining of HeLa cells. (D) Untreated (negative) HeLa cell, (E). LBE treated cells (500 µg/ml), (F). LBE treated cells (1000µg/ml). Arrow indicates nuclear fragmentations.

### Mitochondrial membrane potential assay

Mitochondrial membrane potential, in situ, is an important indicator of mitochondrial function and dysfunction and the loss of mitochondrial transmembrane potential (ΔΨm) leading to damage of mitochondrial membrane is the main step for mitochondrion-dependent apoptotic pathway. HeLa cells were treated with the two optimum concentrations of LBE such as 250 µg/ml in 6 h (Fig.4D, E,F) and in 12 h (Fig.5D, E, F) and 500 µg/ml in 6 h (Fig.4G, H, I) and in 12h (Fig.5G, H,I) *in vitro* with compare to the control (Fig.4A, B, C;) Fig. 5A, B, C) and we examined the effect of LBE on inverted fluorescence microscopy (Olympus, Japan) by JC1 retention as compared with negative control counterparts and it showed that LBE treatment significantly reduced JC1 fluorescence in HeLa cells, suggesting that the mitochondrial pathway is involved in LBE-induced apoptosis. In addition to that we also studied it by flow cytometry (FACS Calibri BD), and the (Fig.6A, B, C, D, E, F, G) showed that LBE disrupted and did dysfunction of mitochondrial transmembrane potential.

**Figure 4.**
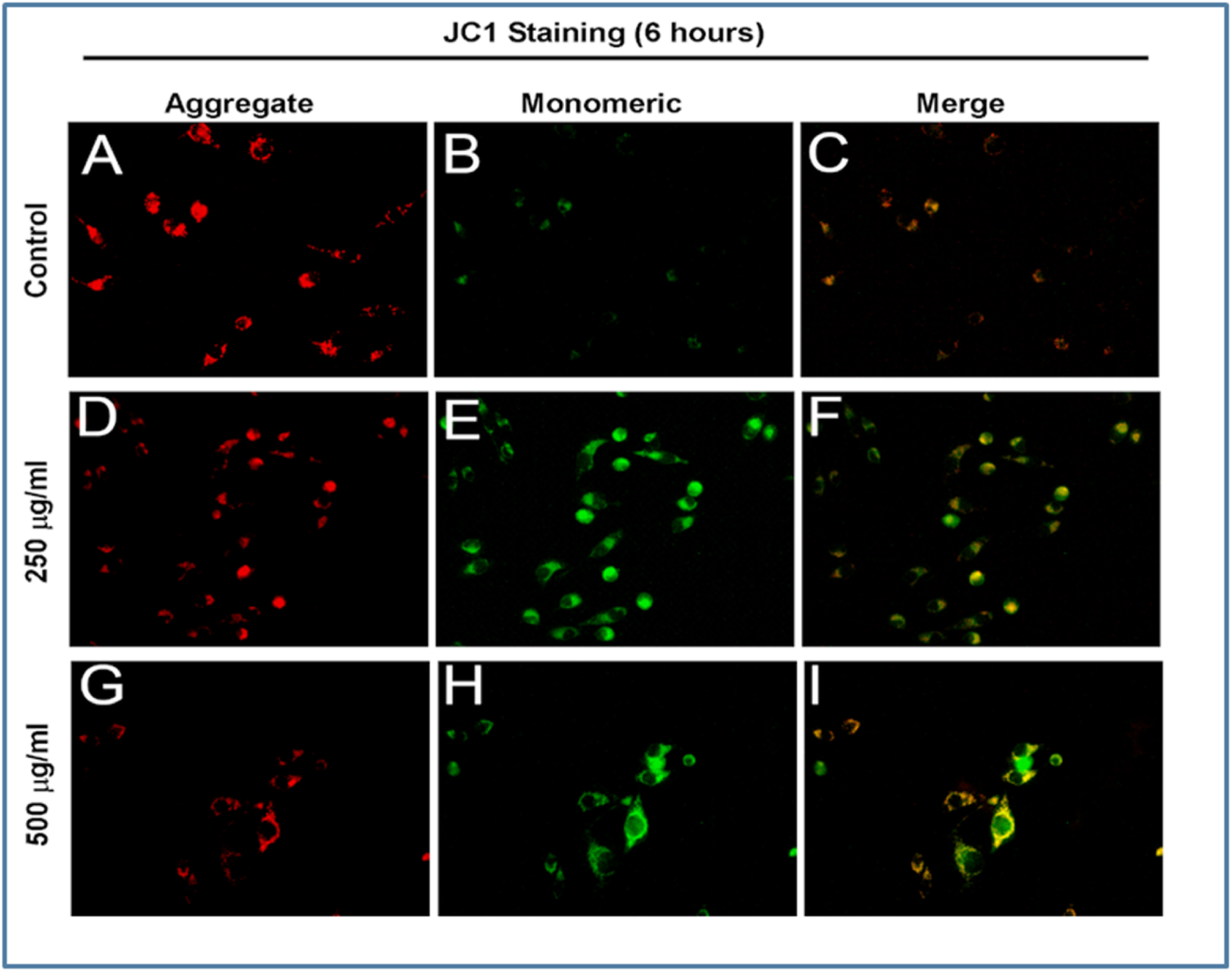
Microscopic image of MMP by JC1 staining in HeLa cell line. Untreated (negative) HeLa cells (A) (aggregate), (B) (monomeric), (C) (merged). Treated cells (250 µg/ml) (D) (aggregate), (E) (monomeric),(F) (merged) and treated cells (500 µg/ml) (G) (aggregate) (H) (monomeric) (I) (merged) Indicate LBE induced time and concentration dependent mitochondrial membrane potential changes (Red to green fluorescence shift) in 6 h and 12 h.

**Figure 5.**
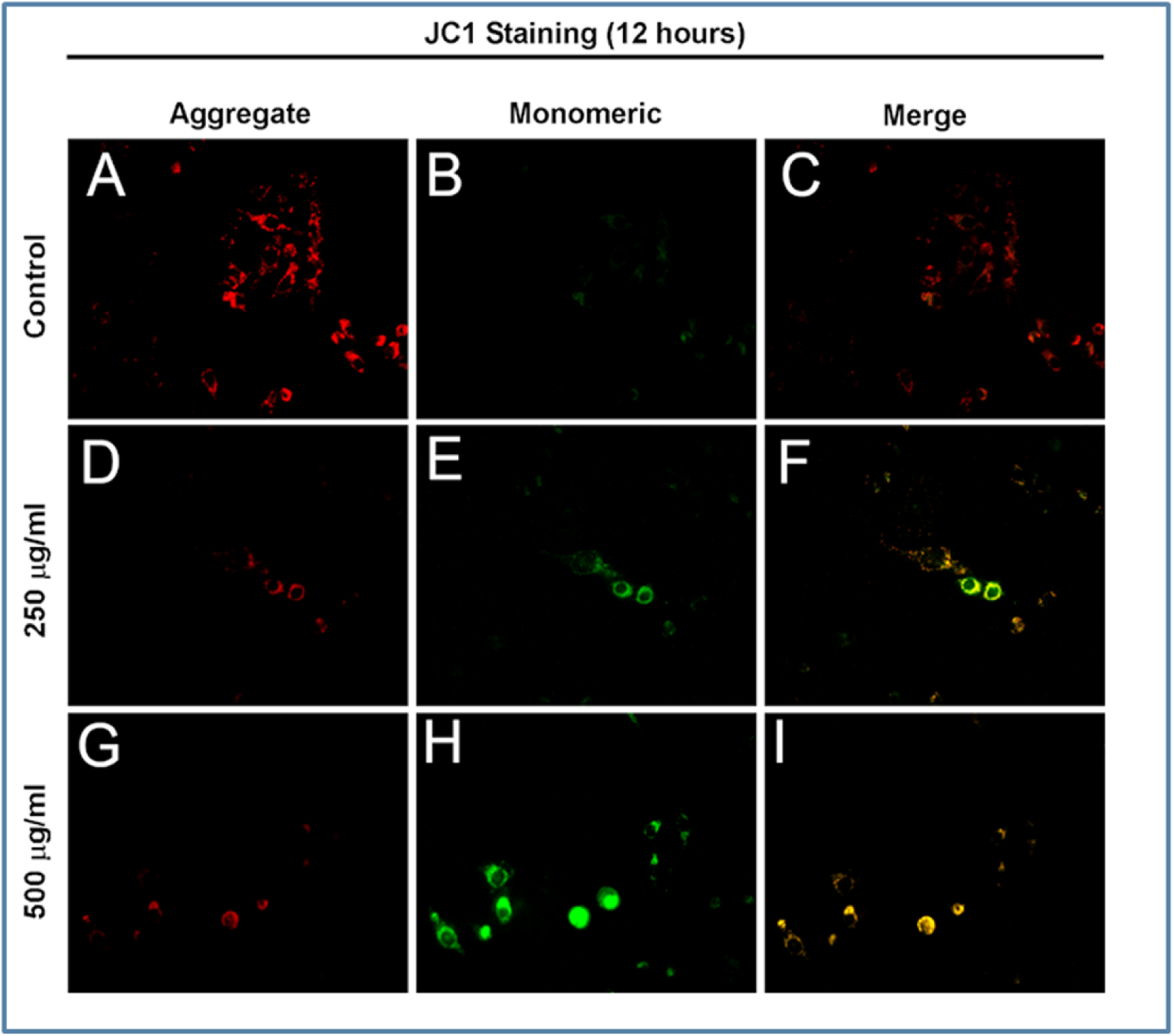
Microscopic image of MMP by JC1 staining in HeLa cell line. Untreated (negative) HeLa cells (A) (aggregate), (B) (monomeric), (C) (merged). Treated cells (250 µg/ml) (D) (aggregate), (E) (monomeric),(F) (merged) and treated cells (500 µg/ml) (G) (aggregate) (H) (monomeric) (I) (merged) Indicate LBE induced time and concentration dependent mitochondrial membrane potential changes (Red to green fluorescence shift) in 6 h and 12 h.

**Figure 6.**
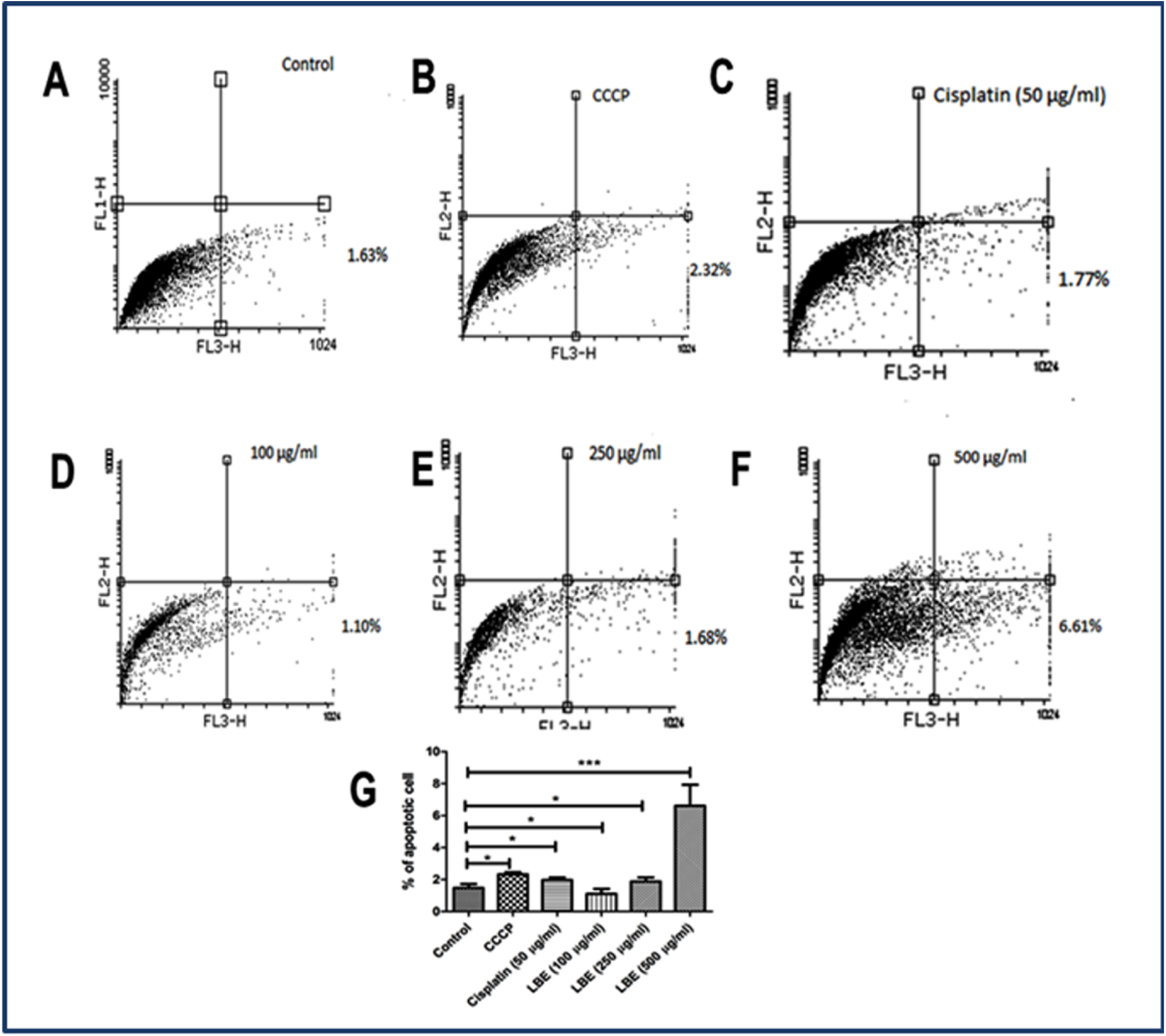
Flowcytometric data on mitochondrial membrane potential changes in HeLa cells. (A) Untreated (negative) cells. (B) CCP treated. (C) Cisplatin (50µg/ml). Treated cells (D, E, F, G), graph (% apoptotic cells). Data are representative as a mean ±SEM of three independent experiments indicates *p<0.05, **p<0.01, ***p<0.001 as compared to control

### Intra cellular ROS generation assay

To investigate the intracellular levels of ROS, the cell-permeable probe H_2_DCFDA was utilized. Non-fluorescent H_2_DCFDA, which is hydrolyzed to DCFH inside the cells, yields highly fluorescent DCFDA in the presence of intracellular H_2_O_2_ and associated peroxides. Whether or not LBE increased or decreased the generation of ROS was subsequently examined. As shown in (Fig.7), LBE treatment (500, 750 µg/ml) increased the generation of ROS in HeLa cells, as determined by monitoring DCF fluorescence, when the dose of LBE was gradually increased. Also we studied ROS by FACS (Fig. 8)

**Figure 7.**
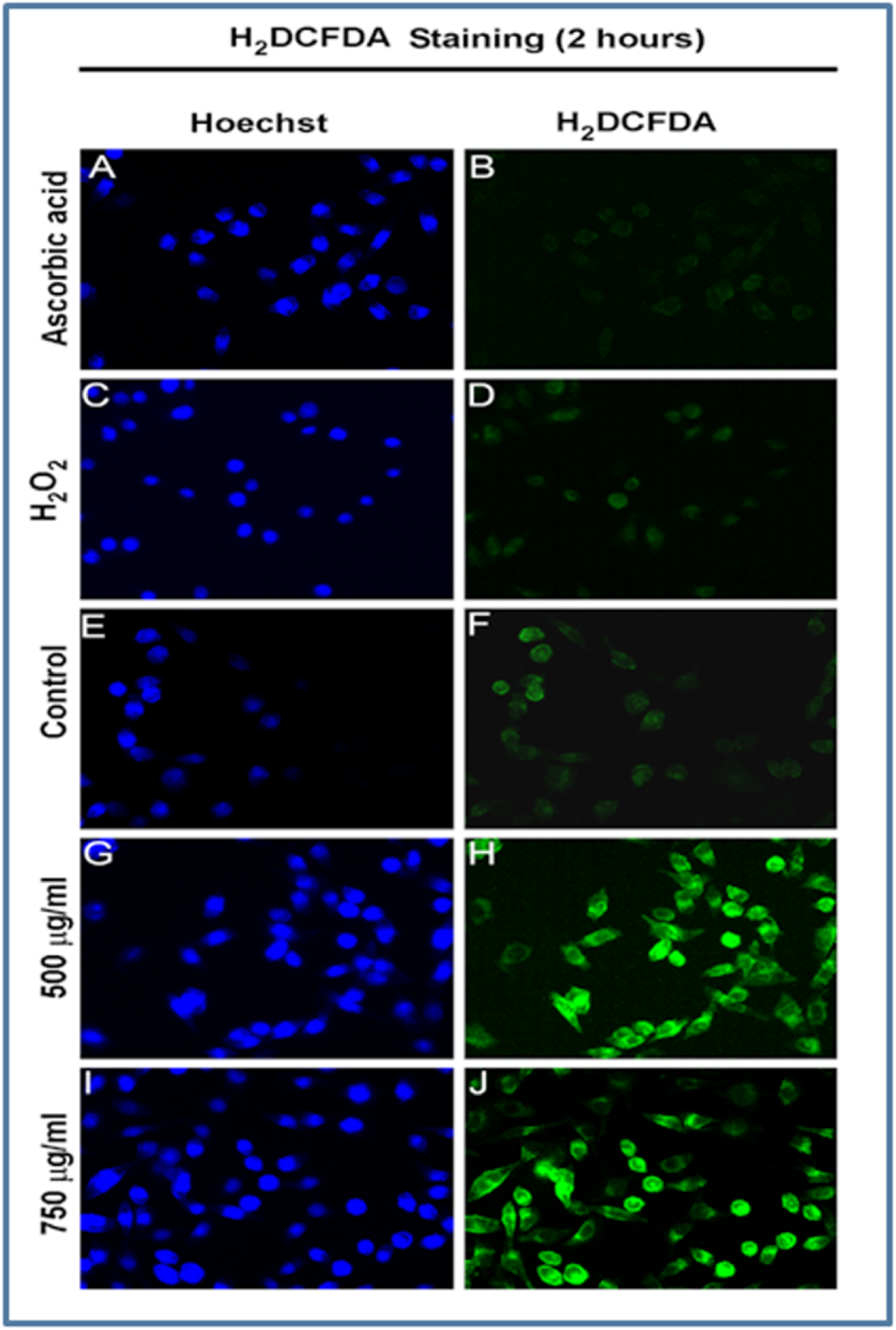
Detection of ROS using carboxy -H_2_DCFDA dye. HeLa cells were treated with H_2_O_2_ were compared to non-treated cells. ROS induces the modification of carboxy-H_2_DCFDA that fluoresces green as detected by flow cytometry, the fluorescent peak in H_2_O_2_ treated cells shift compared to the peaks in controls (H_2_O_2_ treated cells stained with oxidation insensitive dye and non-treated cells stained with carboxy-H_2_DCFDA). Results confirm the presence of ROS in treated cells.

**Figure 8.**
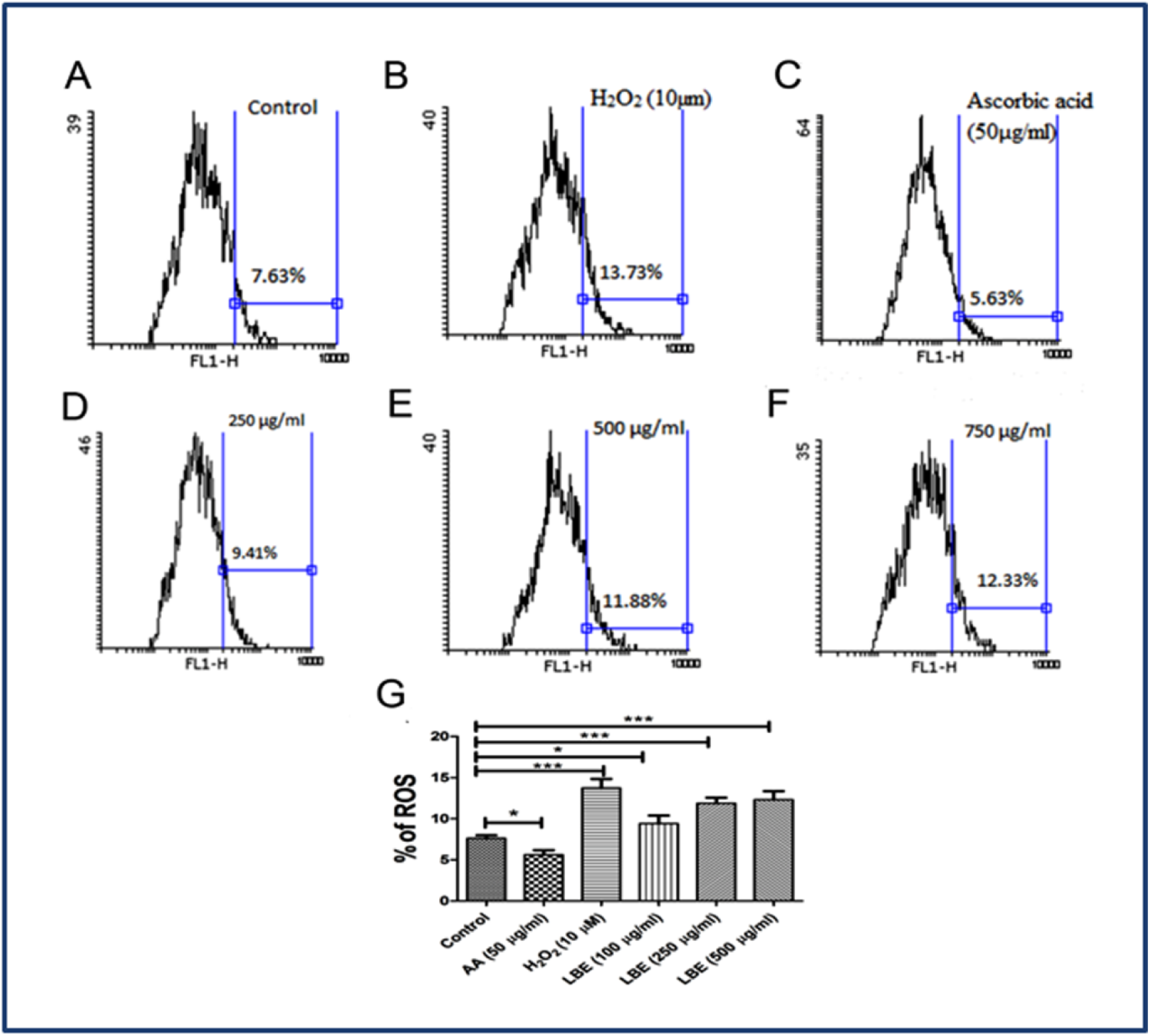
Flow cytometry data of intracellular ROS generation. Untreated (negative) (A), H_2_O_2_ (B), Ascorbic acid (C), treated (D, E, F). Percentage of ROS generation graph (G). Data are representative as a mean ±SEM of three independent experiments indicates *p<0.05, **p<0.01, ***p<0.001 as compared to control

### Cell cycle distribution

We study the cell cycle distribution of HeLa cells in FACS using PI staining. With the treatment of LBE proportion of the cell in the G_2_/M region increased as a dosage dependent and time dependent manner. The G_2_/M population of HeLa cell *in vitro* increase from 16.11 % (control), to 19.55 % (100 µg/ml), 20.21% (250 µg/ml), and 20.62 % (500 µg/ml) respectively in 6 h time duration (Fig.9A, B, C, D, E) In addition to 12 h time duration the HeLa cell was arrested in G_2_/M phase 17.62% (100 µg/ml), 17.64% (250 µg/ml), and 20.04% (500 µg/ml) with compared to the control 16.87% (Fig.10A, B, C, D, E).

**Figure 9.**
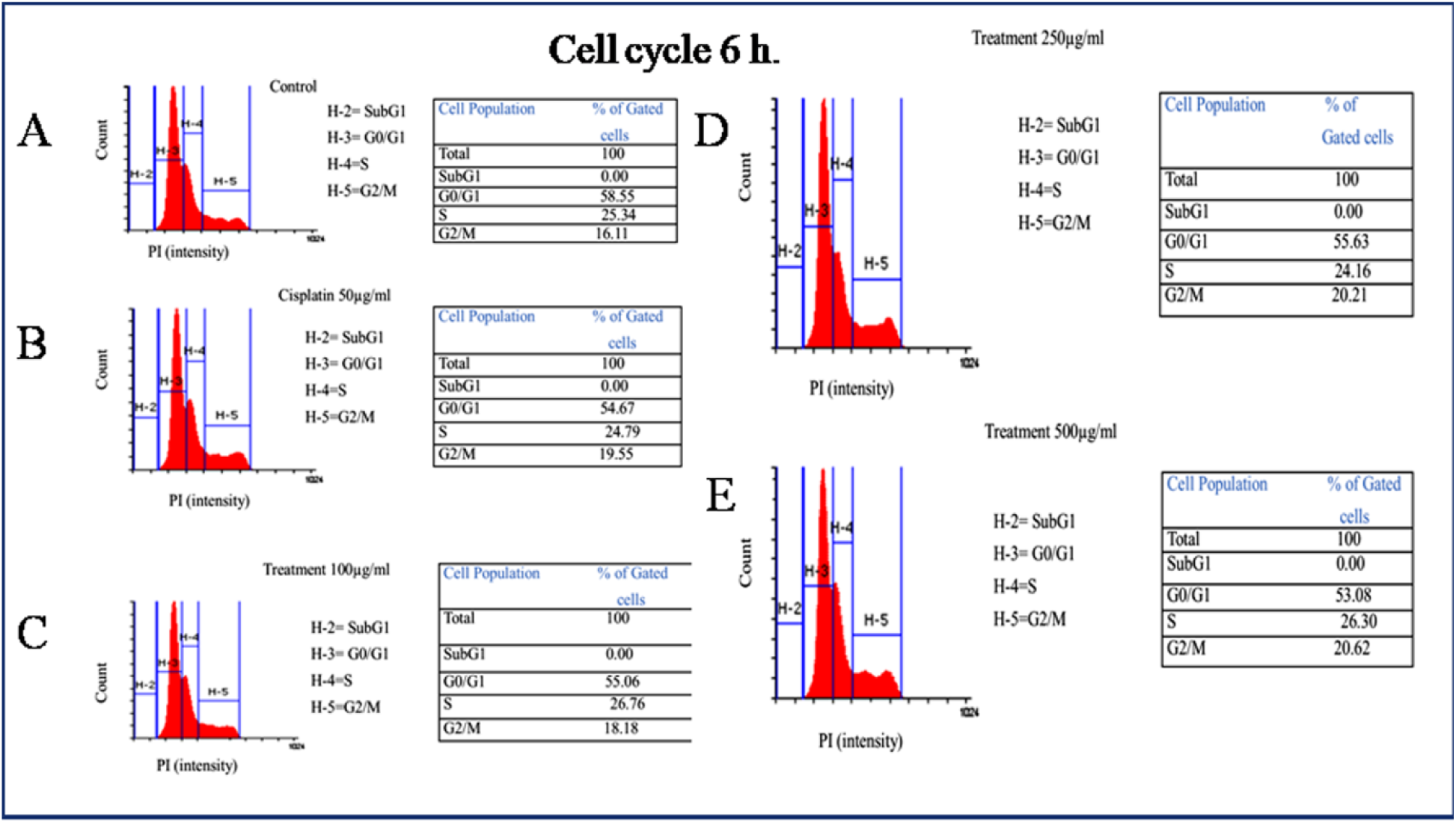
(A) Control (negative).(B, C, D &E) LBE interferes with cell cycle population by inducing G2/M arrest of HeLa cell *in vitro*, in 6 h and 12 h.

**Figure 10.**
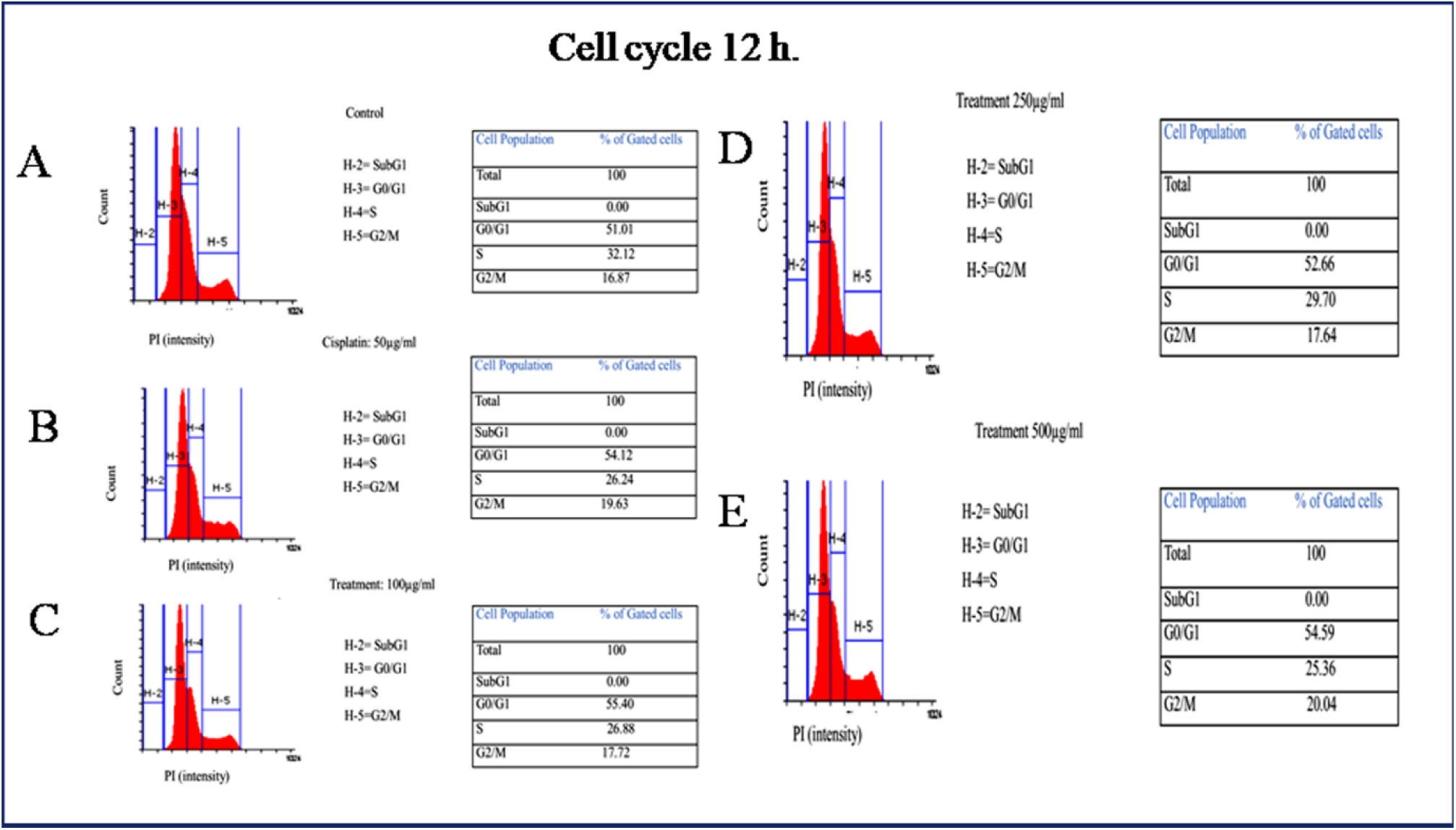
(A) Control (negative).(B, C, D &E) LBE interferes with cell cycle population by inducing G2/M arrest of HeLa cell *in vitro*, in 6 h and 12 h.

### Annexin-V assay

In annexin-V assay four different phenotypes were observed. (1) V-/PI/-(lower left quadrant) indicated as variable cells. (2) V+/PI/-(lower right quadrant), usually estimated as apoptotic cells.(3) V-/PI/+(Upper left quadrant) showed necrotic cells and (4) V+/PI/ +(upper right quadrant) indicated as a late apoptotic cells. The treatment of LBE increased in the population of apoptotic cells was observed with a concomitant decrease in the percentage of viable cells. Compared to the negative control V+/PI/-cells increased 7.09%, 13.5% and 18.1% respectively by the treatment of 250, 500 and 750 µg/ml (Fig.11A, C, D,) respectively, where as in 1000 µg/ml showed V+/PI/+ cells 62.77% in 24 h (Fig 11E).

**Figure 11.**
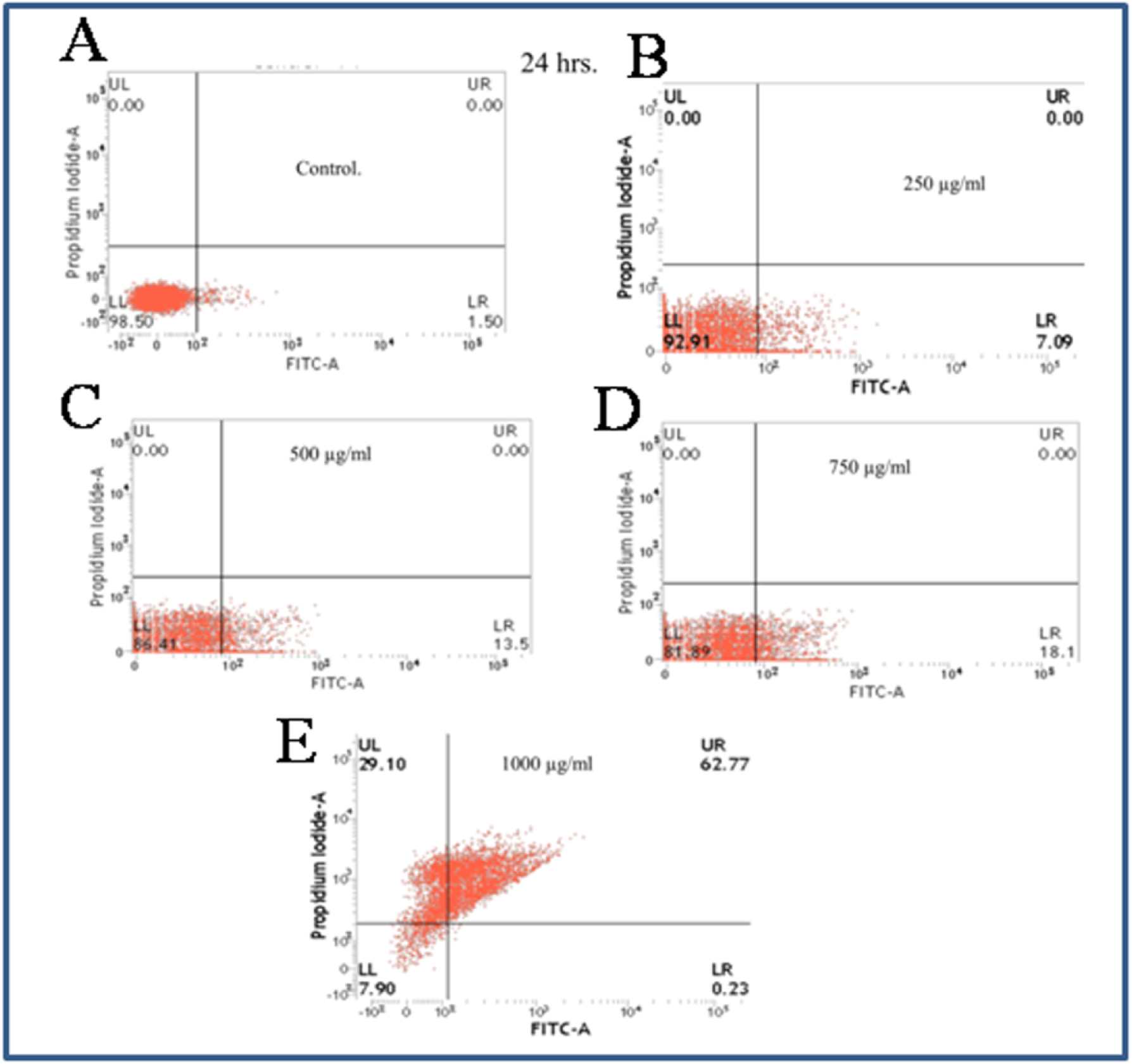
Annexin V Assay in HeLa cells *in vitro* by FACS. (A) Control (Negative) cells. (B, C, D, & E) treated cells. Quadrants: lower left: live cells; lower right: apoptotic cells; upper right: late apoptotic cells; and upper left: necrotic cells.. Data are representative as a mean ±SEM of three independent experiments indicates *p<0.05, **p<0.01, ***p<0.001 as compared to control.

### DNA fragmentation assay

DNA fragmentation of HeLa cells treated with LBE (500,750 and 1000 µg/ml) is easily detected as DNA laddering is prominent in agarose gel electrophoresis (Fig12). Increase of concentration of LBE increased the laddering.

**Figure 12.**
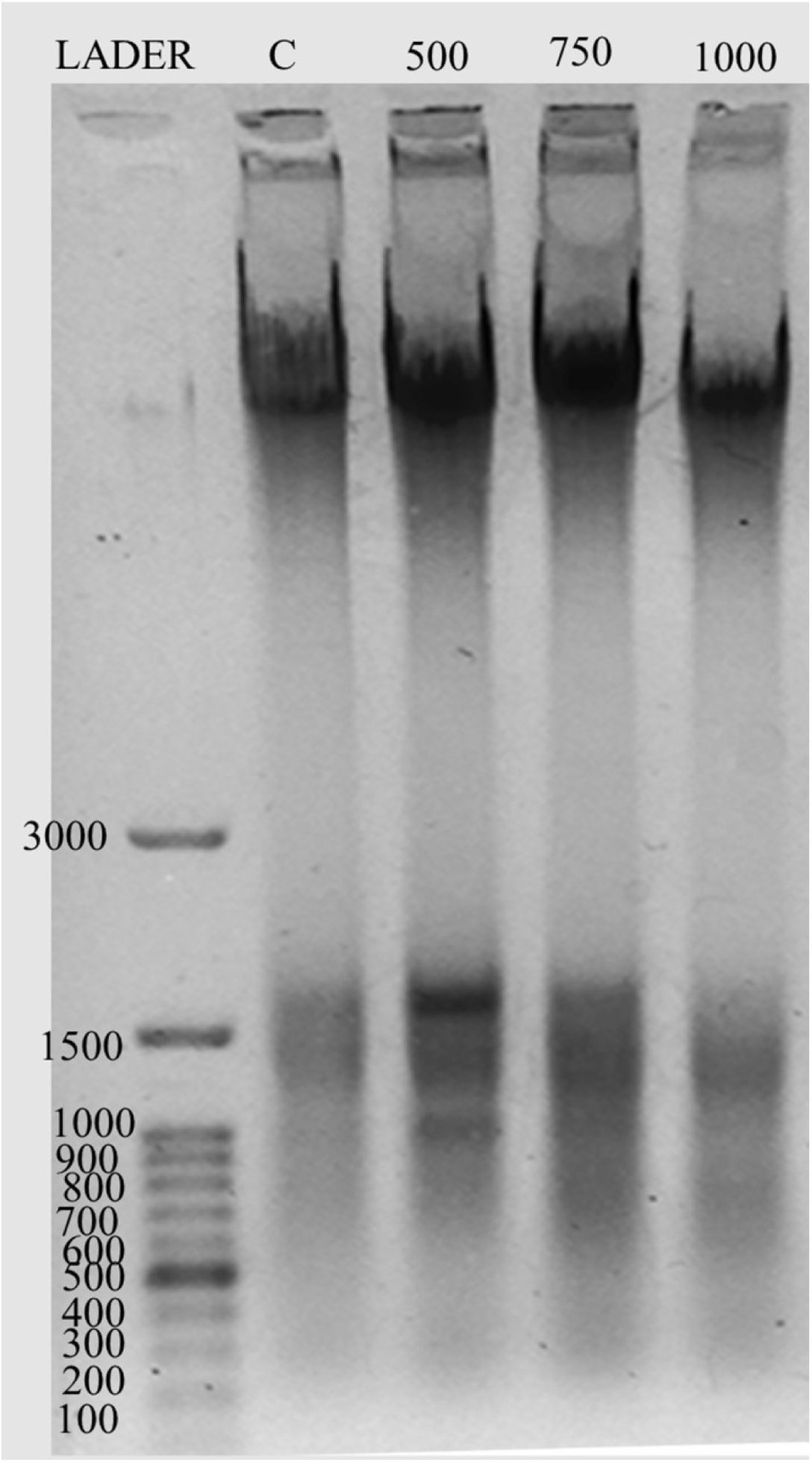
DNA ladder assay. Genomic DNA from HeLa cells was isolated and subjected to electrophoresis in agarose gel (1.8%) and Control (negative) (C) lane showed no fragmented DNA but 1000 μg/ml showed fragmented DNA in the form of ladder after 24 h DNA fragments were visualized under a UV trans-illuminator and compared with a standard marker.

### Induction of Autophagy by LBE

Recently autophagy has gained much attention as the potent alternative mechanism to fight cancer. In fact autophagy is a self-digestive process of cells within auto phagosomes, and is delivered to the lysosome for subsequent degradation and recycling. In our autophagy experiment analyzed by FACS, autophagic cell distribution in negative control was 11.57% (Fig. 13D). It was increased to13.02% (Fig.13F) and 26.76% (Fig.13E) in 250 µg/ml and 500 µg/ml respectively. Our result was also validated by fluorescent microscopic analysis. In negative control, acidic vacuole was not present (Fig.13A), while it was increased with increasing concentrations (Fig.13B, C).

**Figure 13.**
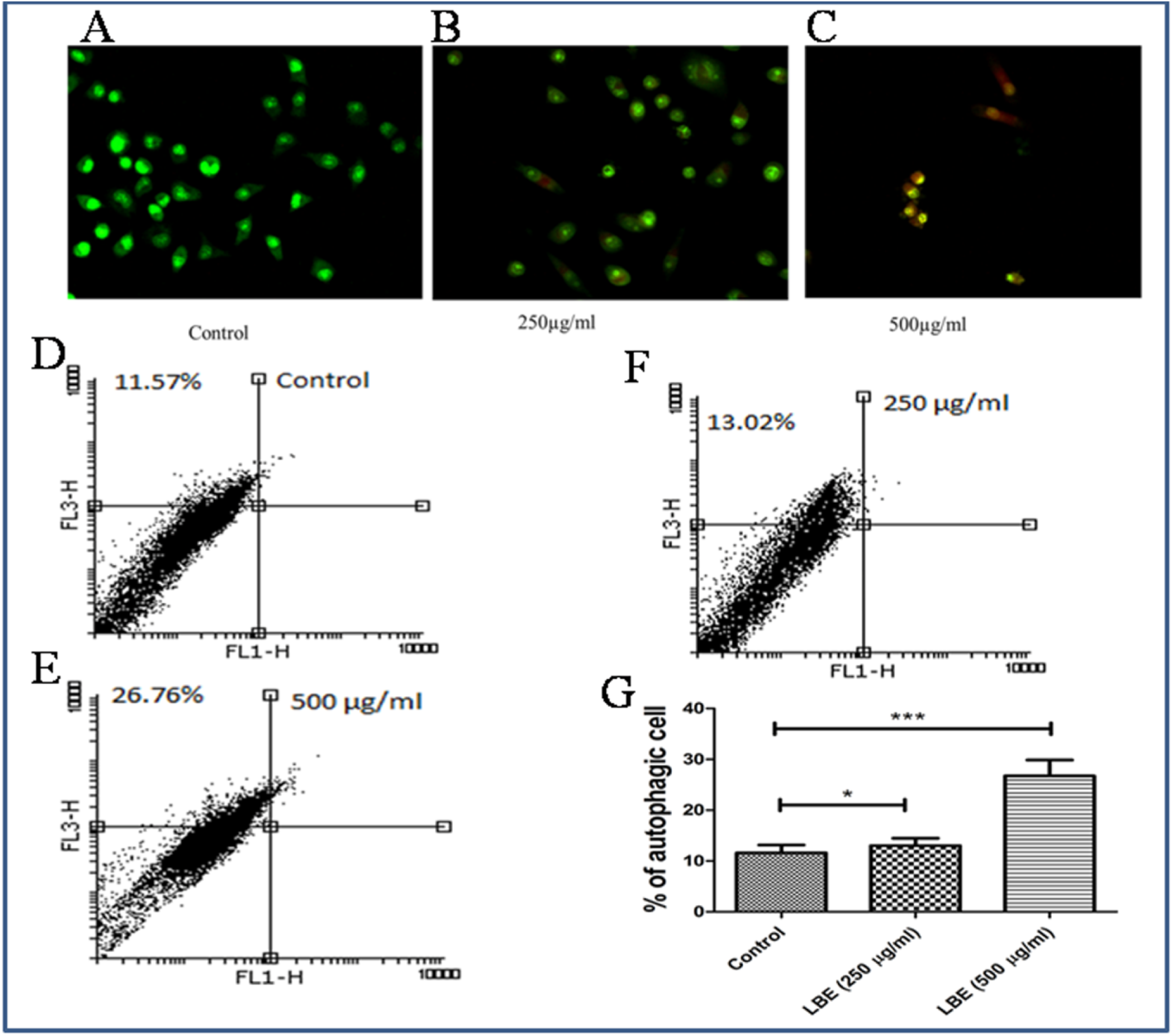
Autophagy of LBE treated in HeLa cell. Fluorescence microscopic image of autophagic cells (A) Negative control, (B &C) treated. (D). Untreated (negative) cells, (E & F). Treated cells, (G).graph (% autophagic cells. Data are representative as a mean ±SEM of three independent experiments indicates *p<0.05, **p<0.01, ***p<0.001 as compared to control.

### F-actin staining

F acting staining under Laser scanning confocal microscope showed that untreated (negative control) cells had no de polymerization in cytoskeleton and DAPI staining also showed the untreated cells are normal and no changes in cytoskeleton. In control (negative) there was no depolymerization of cytoplasmic F-actin. (14B, C). The treated cells with LBE (750 µg/ml) showed that cells are circular and de polymerization of F-actin filaments (14E). Nuclear DAPI staining showed distinct nucleus in case of control (14A), while treated nuclei were condensed and fragmented (14D, F).

**Figure 14.**
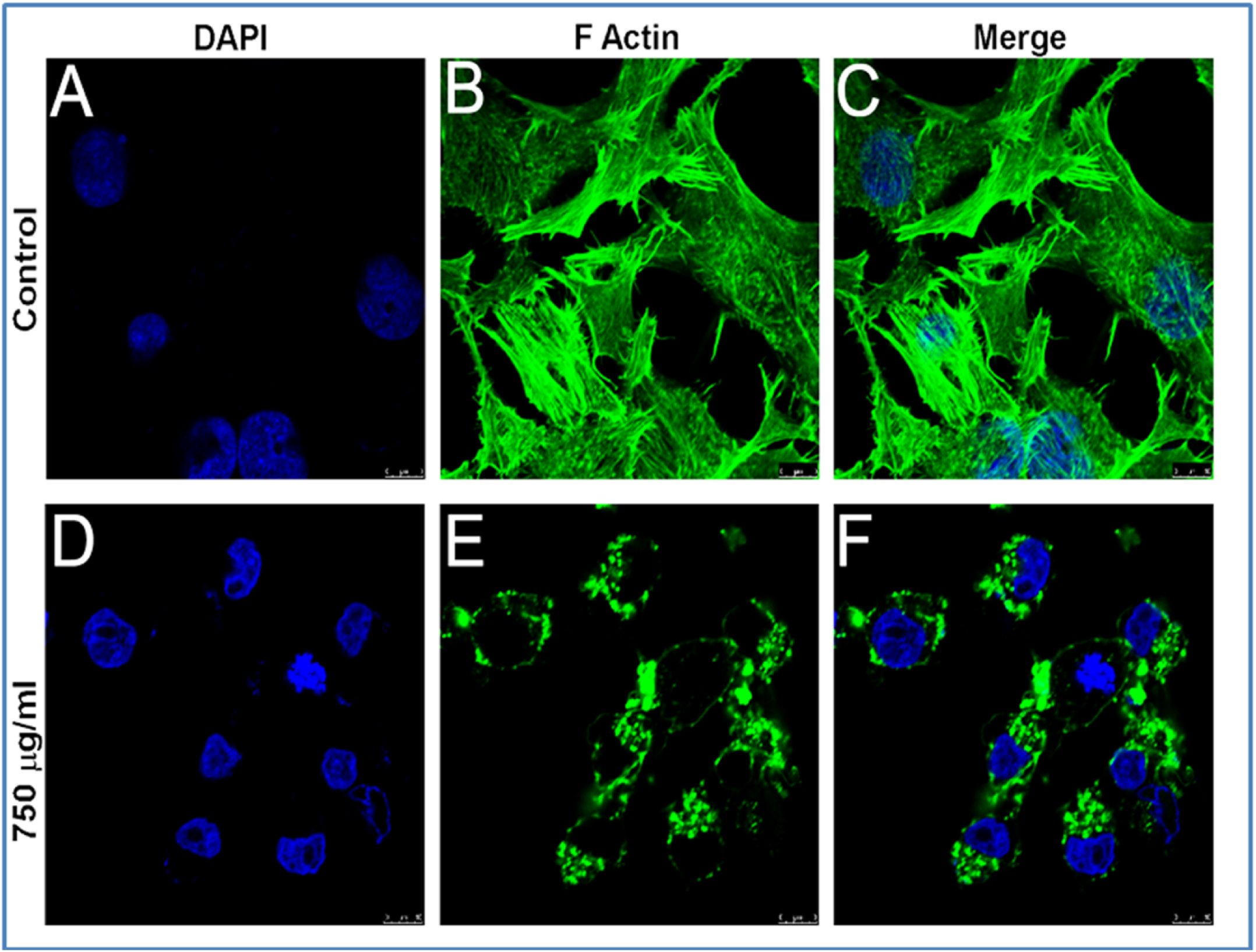
F-actin analysis by confocal microscopy of LBE treated HeLa cell. Control (negative): (A) (DAPI), (B). (F-actin), and (C). (Merged).Treated: (D). (DAPI), (E). (F actin), and (F) (Merged).

### Cancer stem cell population study by Hoechst efflux assay

The Hoechst efflux assay was done by flow cytometer The Side Population(SP) tail contains cells that are able to efflux the Hoechst 33342 dye, The SP of HeLa cells in negative controls was 68.04% (15A), while in LBE treatment, SP of HeLa cells was 46.34% in 250 µg/ml (15B,), 20.81% 500 µg/ml (15C), respectively. Verapamil (positive control) treated HeLa cells showed 2.10% of SP in 50 µg/ml (15D). Percentage of side population was also presented in bar diagram (1E).

**Figure 15.**
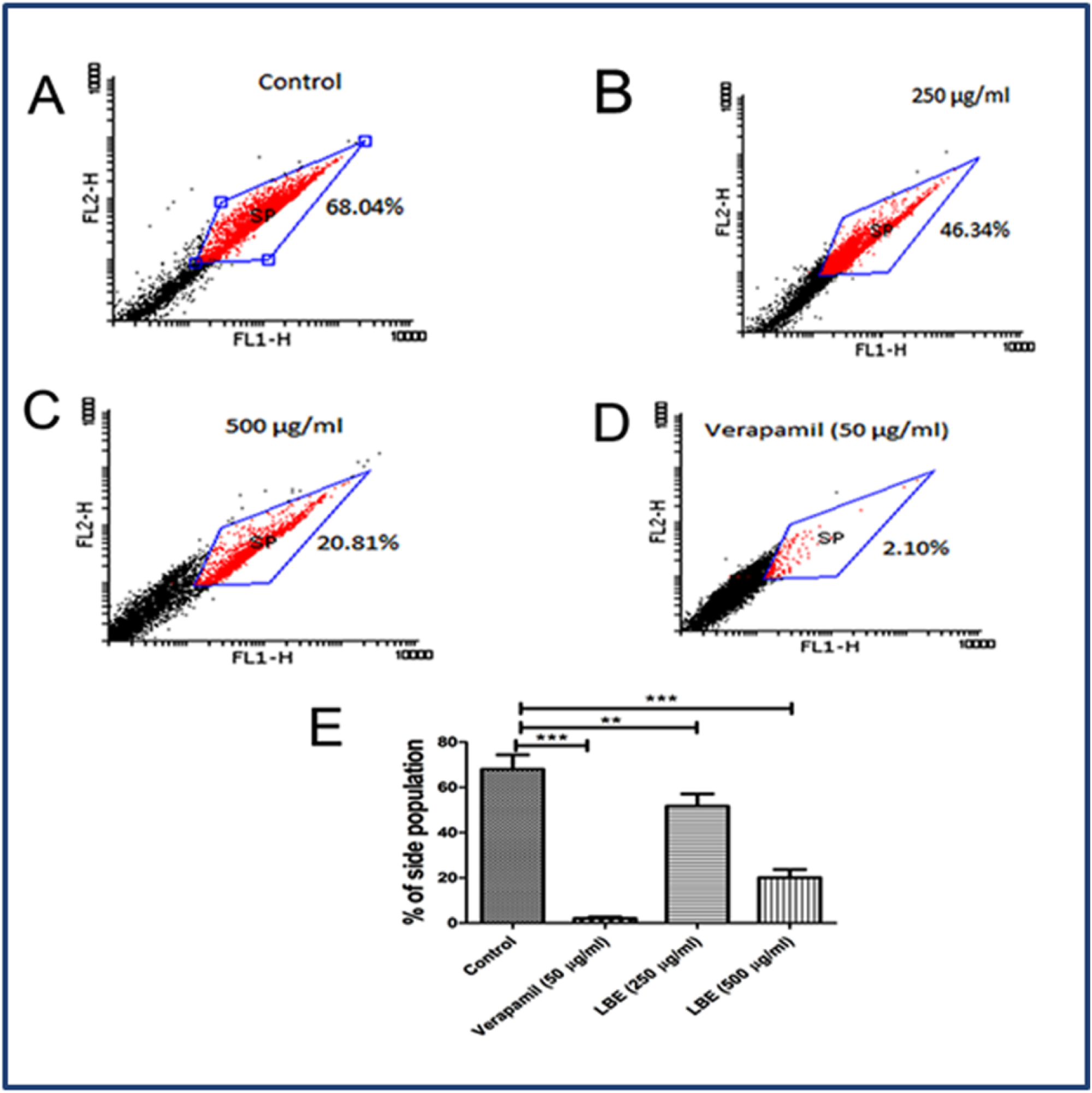
Flow cytometry profiling of stem cell side population. (A). Control (negative), (B&C). Treated (D), Positive control (E). Bar diagram (% of stem cell side population).

### Study of induction of apoptosis by protein expression in Western blotting assay

In order to understand how LBE induced apoptosis of cancer cells and inhibited cell proliferation, the gene expressions of pro- and anti-apoptotic genes such as, pro-caspase 3, pro-Caspase 9, P53, PARP and Bcl-2, were detected. We found that treatment with LBE decreased the expression of gene, pro-caspase 3, pro-caspase 9, Bcl-2 and PARP (Fig.16) in HeLa cells *in vitro*. More over p53 gene, one of important cancer guard genes showed high expression i.e. up regulated by LBE.

**Figure 16.**
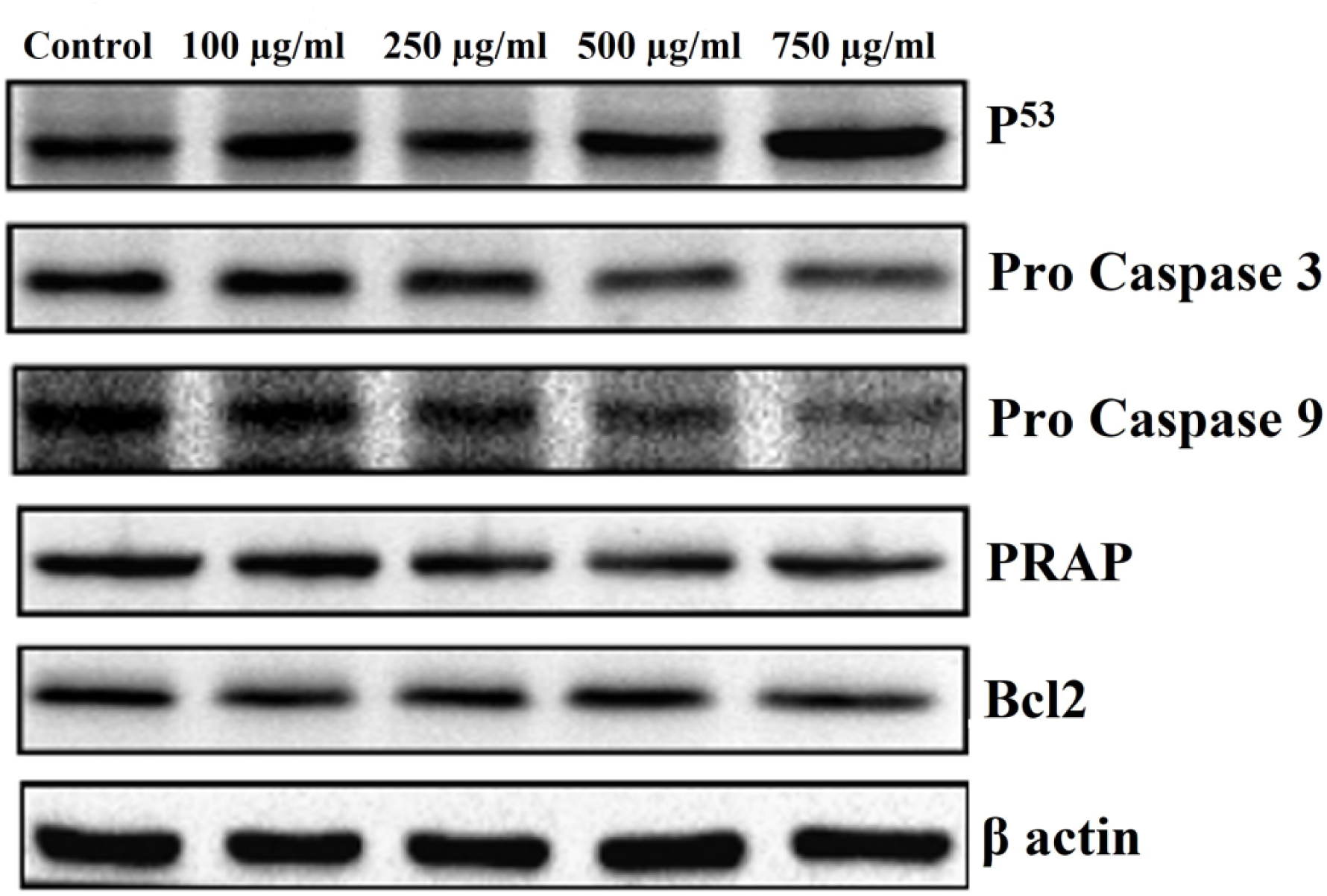
(A) Western immunoblot analysis. Alteration in the expression of p53 pro-caspase 3 pro-caspase 9 BcL-2 and PRAP: Equal loading of protein in the lanes was confirmed by beta actin.

### Anti-migration by LBE against HeLa cells by wound healing /scratch assay

Cell migration was an important property of cancer cells. The effect of LBE on cell migration was examined by a wound healing or scratch assay *in vitro* (Fig 17). Cells were cultured in agarose plates 24 h to come 80% confluency. Cells in the plates were scratched by a 200 µl pipette tip in the center of the well and exposed with various concentrations of LBE. Briefly, 50×10^3^ HeLa cell per well were cultured in 24 well plate. After reaching 90% confluency, the center of the culture dishes was scratched with 200 µl pipette tip. Then the cells were washed frequently by using PBS and incubated with LBE at concentration of 0, 500 and 1000 µg/ml. After 24 h image of the cells was taken under 10X magnification.

**Figure 17.**
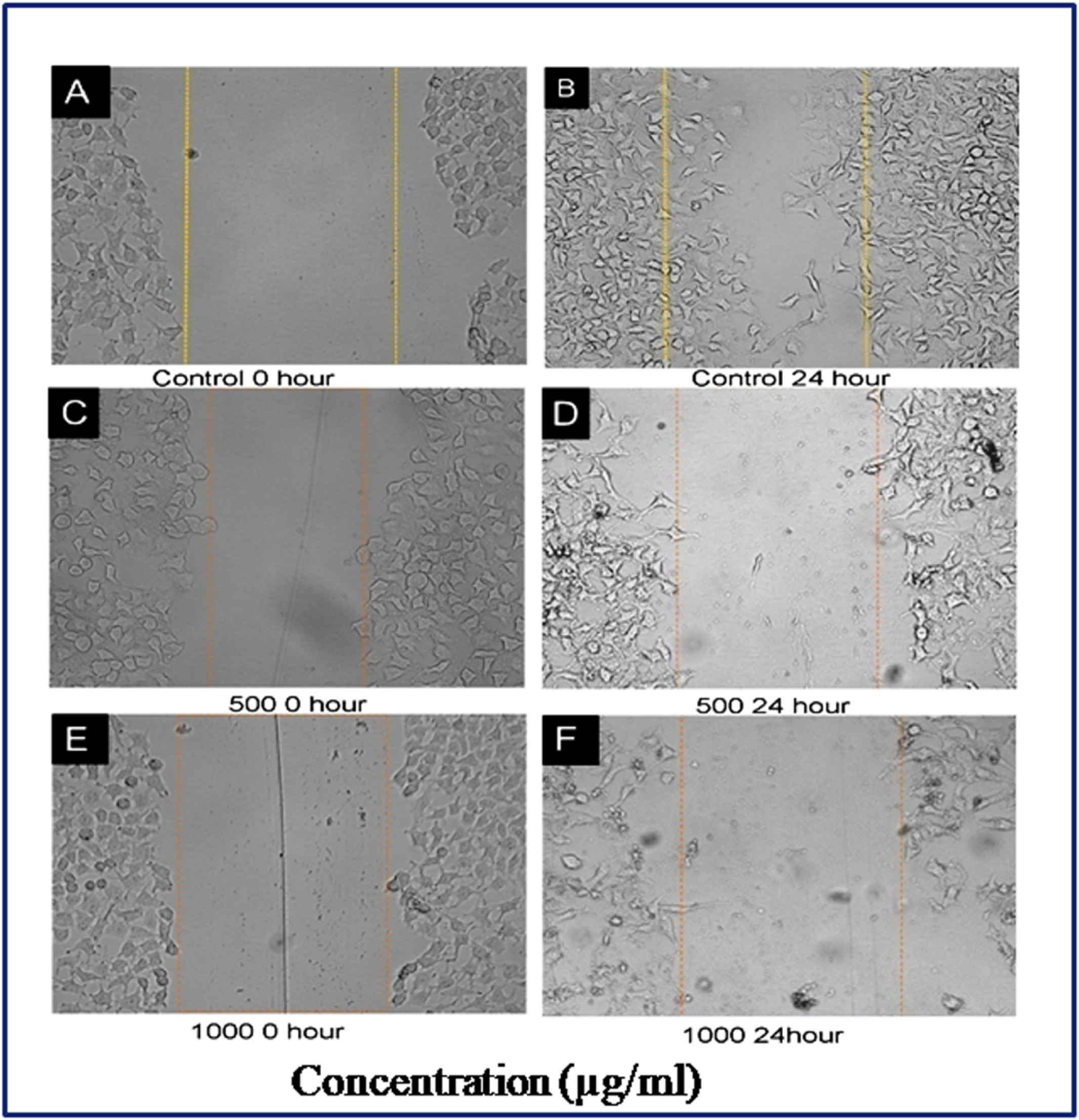
Effect LBE on migration of HeLa Cells in scratch method. (A) No cell migration in control (negative) at 0 h. (B) Cell migration in control at 24 h. Treated cell 0 h (C, E) and 24 h (D, F).

### Cologenic assay

We tried to examine the capacity of the LBE to inhibit the colony formation of HeLa cells on agarose plate. The results showed maximum colony formation of HeLa cells in control (negative) (18A) where as in treatment, LBE decreased the colony formation of this cancer cells gradually when we increased doses from 250 µg/ml to 1250 µg/ml (18 B, C, D, E & F) and in maximum dose 1250 µg/ml (18 F) no cells were found in 24 h. Treatment. The property of anchorage-independent growth of cancer cells, *in vitro* is one of the important properties, and colony formation is correlated with the *in vivo* oncogenic potential of cancer cells.

**Figure 18.**
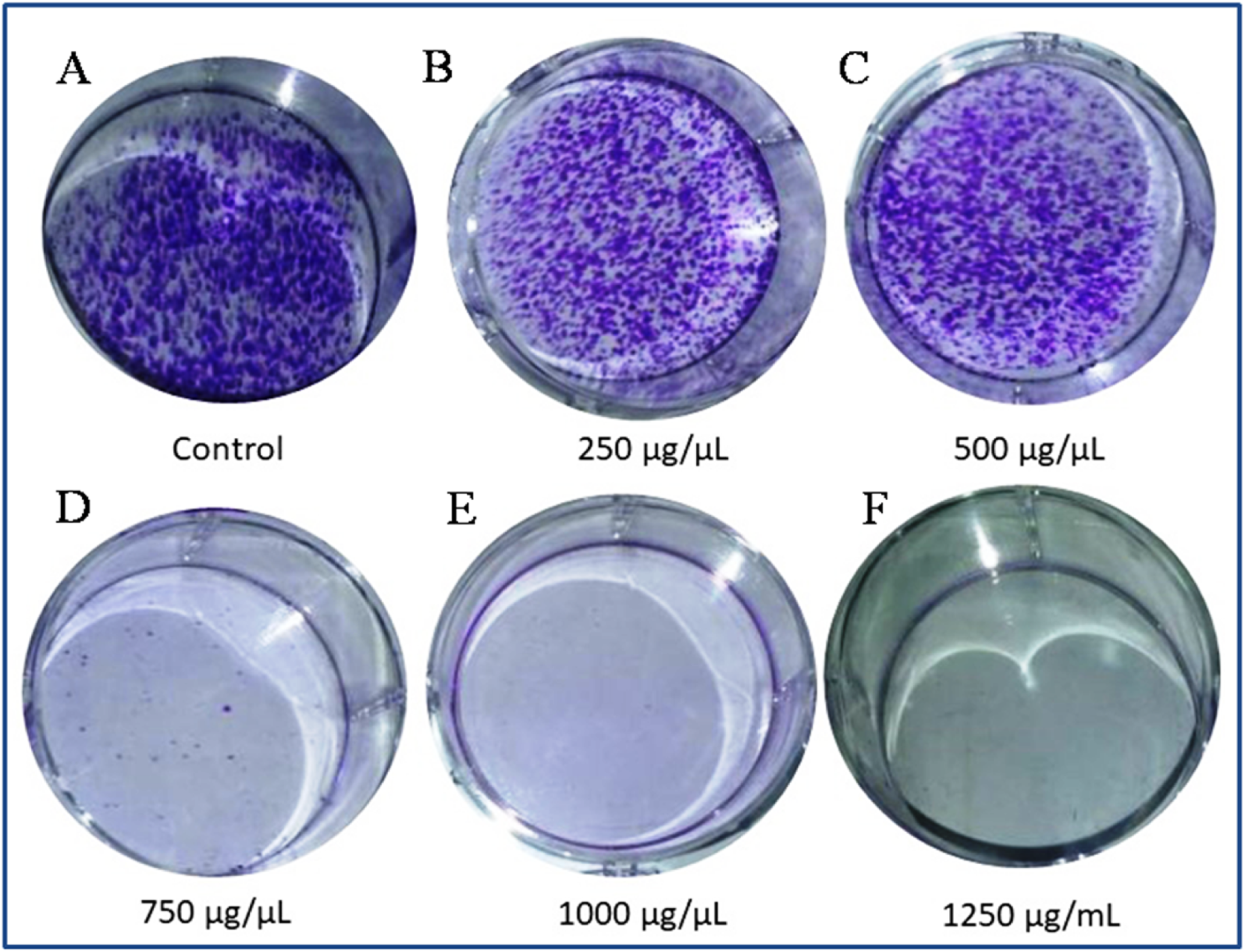
Effect of LBE on colony formation of HeLa cells *in vitro*. (A) Control (negative). Treated (B, C, D, E, and F): dose dependent inhibition of clonogenicity in HeLa cell

### *Antitumor effects of LBE treatment in HeLa-implanted mic*e

Growth of the tumors in HeLa cell-implanted mice is shown in (Fig19), and tumor growth (volume) in 50 mg/kg LBE treatment groups were plotted in line plot (Fig.20). Tumors in the HeLa + vehicle group increased in volume after 7 days (0.65±0.10 cm^3^), reaching a maximum of 4.10± 1.59 cm^3^ after 28 days. The tumor volume for the 50 mg/kg LBE treatment groups on days 11, 15, 21, and 28 were 0.49± 0.07, 0.23 ±0.05, 0.14±0.03, and 0.04 ± 0.01cm^3^, respectively (Table 1). When the weight of the tumors in this group was measured on day 28, following the sacrifice of the mice, the tumors were of the minimum weight (214 ±12.34 mg), whereas, in the HeLa + vehicle control group (negative), the tumors attained the maximum volume (4.10±1.59 cm^3^). Treatment with 50 mg LBE/kg of body weight of mice led to a marked reduction in the volume (93.22±9.2% cm^3^) and weight (90.42 ±9.55 %) of the tumors. However, the reduction of tumor volume in the HeLa + LBE 10 treatment group (3.95 ± 2.00 cm3) did not reach a significant level compared with the HeLa+ vehicle group (4.10± 1.59 cm^3^). The visceral organs (kidney, liver, spleen, and lungs) were morphologically observed, and they were revealed to be normal and intact: No disorganization, no lesions and no abnormal growth were identified.

**Figure 19.**
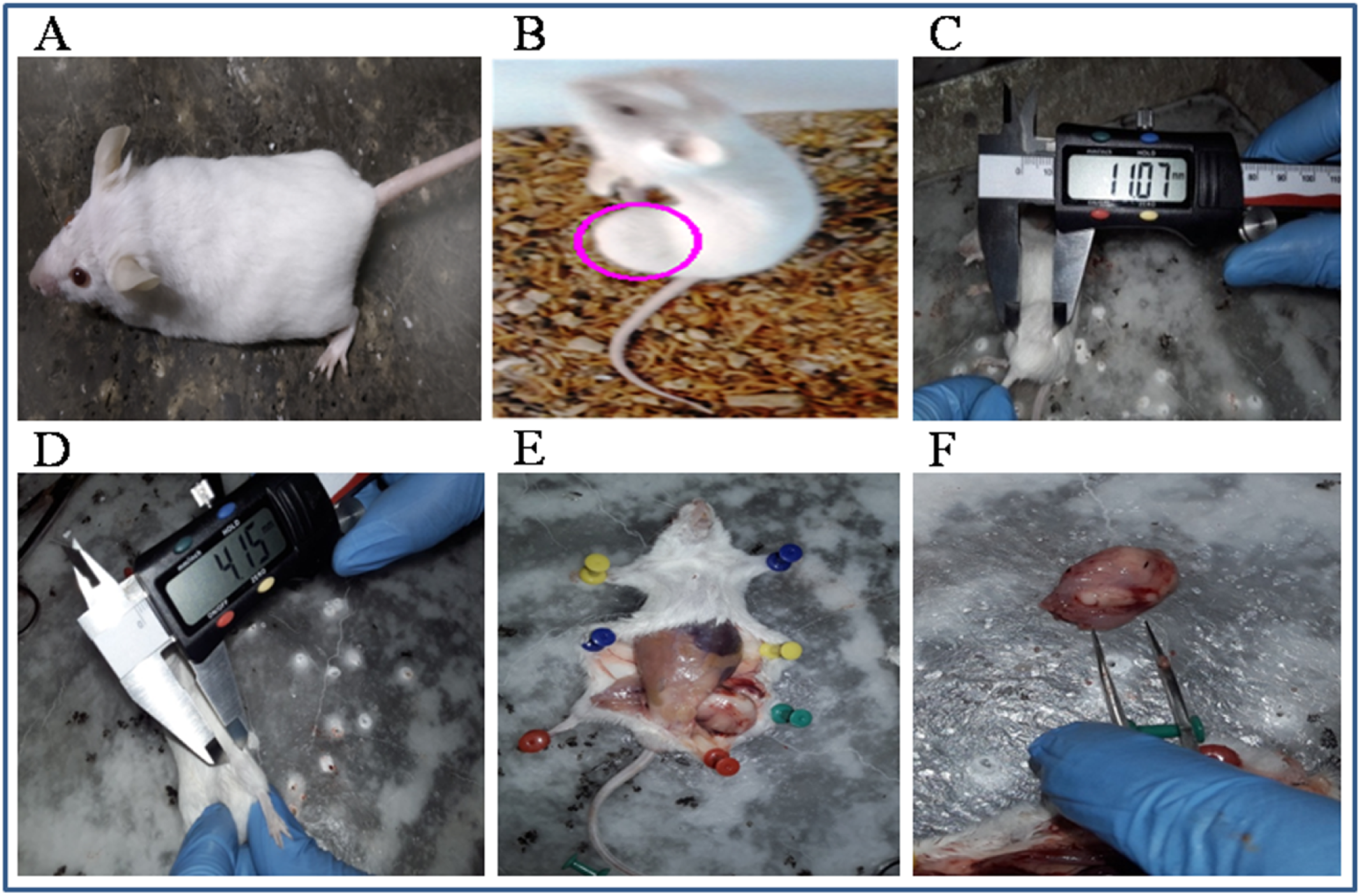
Effects of LBE on tumor growth on HeLa implanted mice. LBE suppresses the growth of cervical cancer tumors in swiss albino mice. Mice were inoculated subcutaneously in the right flank with 2×10^6^ HeLa cells. Tumor volume was measured twice/week using a caliper and calculated as (width)^2^ × length/2. Representative images were captured at the end of therapy, (A) shows control mice (B) Tumor bearing mice, (C) measurement of tumor treated (LBE 10 mg/kg body weight), (D) measurement of tumor treated (LBE 50 mg/kg body weight), (E) sacrificed mouse (dissected), (F) tumor dissection from sacrificed mice.

**Figure 20.**
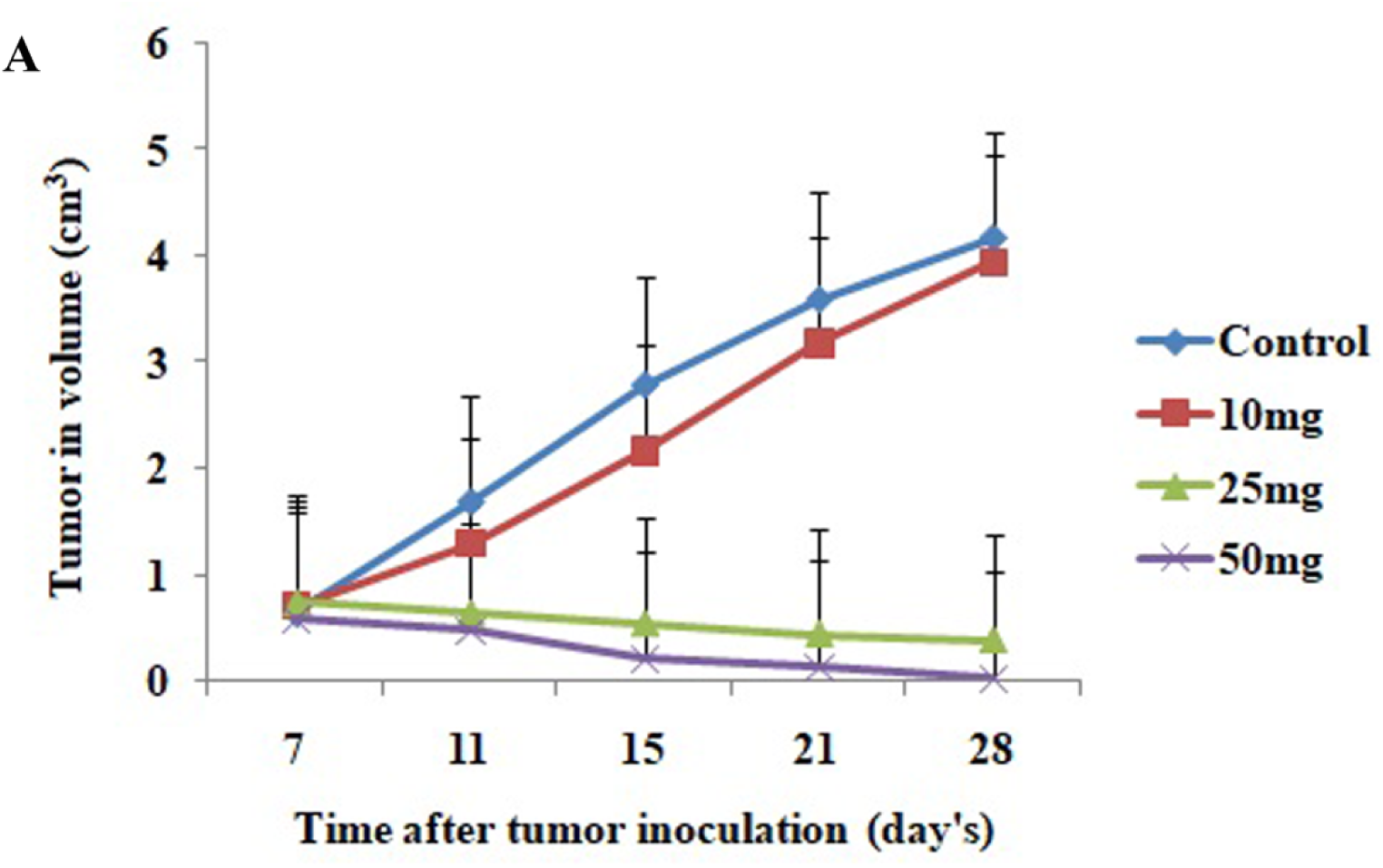
Graphical representation (line plot) of the effects of LBE on tumor growth (volume) *in vivo.* mg/kg indicates mg LBE per kg body weight of mice. The data represented as mean ± SD for the three different experiments performed in triplicate. Error bars of ± SD are inserted in figure

**Table 1.** Tumor progression and reduction in volume and percentage of reduction at various doses (mg/Kg body weight of mice) of LBE

| Treatment No. of ( Day) | Tumor volume in cm<sup>3</sup>± SD |  |  |  |
| --- | --- | --- | --- | --- |
|  | HeLa+vehicle Group | 10 mg LBE /Kg Group | 25mg LBE /Kg Group | 50mg LBE /Kg Group |
| 1st(after 7 days) | 0.65±0.10 | 0.71± 0.12 | 0.75 ±0.09<br>*(0.0) | 0.59±0.09<br>(0.0) |
| 2<sup>nd</sup> (11<sup>th</sup> Day) | 2.03±0.15 | 0.97 ±0.15 | 0.65 ±0.10<br>( 13.33± 2.12) | 0.49± 0.07<br>(16.94± 2.52) |
| 3<sup>rd</sup>(15<sup>th</sup> Day) | 3.02± 1.00 | 2.17 ±1.15 | 0.55±0.10<br>(26.66± 3.23) | 0.23 ±0.05<br>(61.01± 4.34) |
| 4<sup>th</sup> (21<sup>st</sup> Day) | 3.57 ± 1.30 | 3.19 ± 1.00 | 0.45±0.05<br>(40±3.90) | 0.14±0.03<br>(76.27± 6.12) |
| 5<sup>th</sup> (28<sup>th</sup> Day) | 4.10± 1.59 | 3.95 ± 2.00 | 0.39 ± 0.90<br>(50± 4.00) | 0.04 ± 0.01<br>(93.22±9.2) |

Note: \*Data within parentheses indicate percentage (%) of tumor reduction or regression in volume. The calculation of percentage of reduction of tumor volume was done with respect to the tumor volume generated after 7 days of implantation of HeLa in 25mg LBE /Kg Group and 50mg LBE /Kg Group.

### Gas chromatography-mass spectrometry (GC-MS) analysis

GC-MS analysis is powerful tool for qualitative and quantitative analysis of various compounds present in natural products. And widely being applied in medical, biological and food research ^34-37^. In present study the GC-MS analysis of the LBE of *L. betulina* from the NIST inbuilt, Library Data Bank showed the presence of 69 compounds (Fig 21). Of the 69 compounds obtained, 32 compounds had the higher matching probability above 50% with the standard compounds of the NIST Library data Bank (Table 1). Out of which, 14 compounds found to be in higher proportion with the percentage peak area >1. Two of the 14 compounds 9,12-Octadecadienoic acid (Z,Z)(Mwt :280.452 g/mol, Chemical formula :<u>C_18_H_32_O_2_</u>) and Ergosta-5,8,22-trien-3-ol, (3.beta22E) (Mwt: 396.6484 and Chemical formula: C_28_H_44_O). Were in a significantly higher proportion with the percentage peak area 22.13 and 19.72 respectively.  Next compounds with their % peak area are Tetradecanoic acid (9.87, saturated fatty acid), Linoleic acid ethyl ester (6.43, polyunsaturated), Oleic acid (4.37, 2.42, monounsaturated fatty acid); Hexadecanoic acid, 2-hydroxy-1-(hydroxymethyl) ethylester (2.31 Cyclooctacosane (1.38, saturated fatty acid); Phenol, 2,4-bis (1,1-dimethylethyl) (1.50), Phenol, 2,4,6-tris (16icycle[2.2.1]hept-2-yl), and 4’-Hydroxy-6-methoxyaurone (1.34). Lupeol, which is also present in LBE, is a natural pentacyclic triterpenoid (Table2).

**Figure21.**
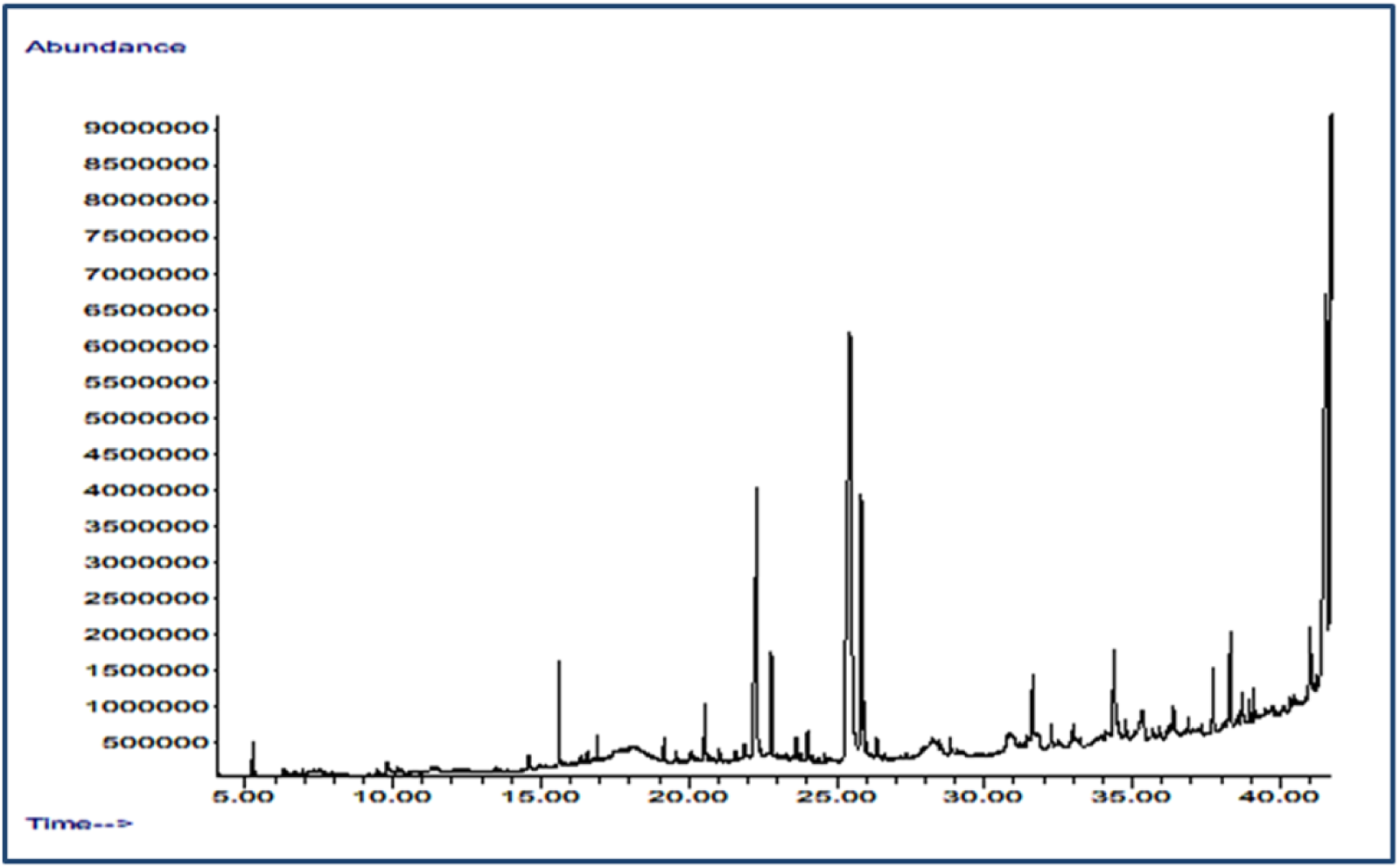
GC-MS chromatogram

**Table 2.** Compounds with quality /probability greater than 50 %, their retention time (RT) and % of peak area in GS-MS

| No | RT | Quality/Probablity | % of peak Area | Compound |
| --- | --- | --- | --- | --- |
| 1 | 5.254 | 74 | 0.27 | Hexylene Glycol |
| 2 | 14.589 | 56 | 0.47 | 2-(2-Hydroxyethyl)piperidine |
| 3 | 15.616 | 94 | 1.50 | Phenol, 2,4-bis(1,1-dimethylethyl) |
| 4 | 16.342 | 50 | 0.13 | Dodecanoic acid |
| 5 | 16.898 | 95 | 0.34 | Diethyl Phthalate |
| 6 | 19.160 | 96 | 0.39 | Tetradecanoic acid |
| 7 | 19.570 | 97 | 0.22 | 5-Octadecene, (E)- |
| 8 | 20.545 | 93 | 1.23 | Pentadecanoic acid |
| 9 | 21.038 | 78 | 0.22 | Tetradecanoic acid, ethyl ester |
| 10 | 21.578 | 78 | 0.35 | Hexadecanoic acid, methyl ester |
| 11 | 21.889 | 58 | 0.40 | 4-Undecene, (E) |
| 12 | 22.304 | 97 | 9.87 | Tetradecanoic acid |
| 13 | 22.792 | 98 | 2.00 | Hexadecanoic acid, ethyl ester |
| 14 | 23.638 | 76 | 0.50 | Hexadecadienoic acid, methyl ester |
| 15 | 24.016 | 93 | 0.59 | Heptadecanoic acid |
| 16 | 24.592 | 81 | 0.22 | Heptadecanoic acid, ethyl ester |
| 17 | 25.459 | 98 | 22.13 | 9,12-Octadecadienoic acid (Z,Z) |
| 18 | 25.516 | 93 | 4.37 | Oleic Acid |
| 19 | 25.563 | 92 | 2.42 | Oleic Acid |
| 20 | 25.832 | 99 | 6.43 | Linoleic acid ethyl ester |
| 21 | 25.921 | 95 | 1.21 | Ethyl Oleate |
| 22 | 26.367 | 96 | 0.38 | Octadecanoic acid, ethyl ester |
| 23 | 31.628 | 90 | 2.31 | Hexadecanoic acid, 2-hydroxy-1-(hydroxymethyl)ethyl ester |
| 24 | 32.266 | 80 | 0.58 | 1,2-Benzenedicarboxylic acid, diisooctyl ester |
| 25 | 34.373 | 97 | 2.86 | 9,12-Octadecadienoic acid (Z,Z) |
| 26 | 34.773 | 64 | 0.50 | Octadecanoic acid, 2-hydroxy-1-(hydroxymethyl)ethyl ester |
| 27 | 35.333 | 55 | 01.34 | 4'-Hydroxy-6-methoxyaurone |
| 28 | 36.391 | 70 | 0.59 | 2,6,10-Dodecatrien-1-ol, 3,7,11-tr |
| 29 | 37.725 | 78 | 1.38 | Cyclooctacosane |
| 30 | 38.711 | 52 | 0.90 | Phenol, 2,4,6-tris(17icyclo[2.2.1]hept-2-yl)- |
| 31 | 41.134 | 90 | 0.23 | Stannane, dibutylcyclohexylmethyl |
| 32 | 41.549 | 91 | 19.72 | Ergosta-5,8,22-trien-3-ol,<br>(3.beta.22E) |

## DISCUSSION

The current findings demonstrated that LBE of *L. betulina* is very potent killer of HeLa, CaSki and SiHa as validated by MTT assay, and present elaborate experimental works on various parameters such as inhibition of proliferation, induction of apoptosis, reduction of mitochondrial membrane potential, generation of intracellular ROS, induction of autophagy of HeLa cell line, de-polymerization of F-actin of cytoplasm, cancer stem cell population study and anti-metastatic assay (migration and cologenic assay) were not conducted by any other worker but previously extracts of different mushrooms on other cell lines were reported by other scientists. As per the report of Liu *et al*.^36^, the EE of *L.betulina* displayed strong anticancer activity against cancer cell line MDA-MB-231, with IC_50_ value of 51.46 μg/ml, and there was 83.15% inhibition at a concentration of 200 μg/ml (MTT assay). Before that Oyetayo^8^ evaluated the hex-anolic (LHE), ethyl acetate (LEA) and ethanol (LET) extracts of *L .betulina* for cytotoxic effect (%) against some cancer cell lines such as U251 (glioblastoma), PC-3 (prostatic adenocarcinoma), K562 (chronic myelogenous leukemia), HCT-15 (colorectal adenocarcinoma), MCF-7 (breast cancer cell line) and SKLU-1 (lung adenocarcinoma). In our experiment of MTT assay against HeLa cells, LBE showed 70% inhibition and IC_50_ was 492.52±0.56µ g/ml. In our previous experiments ^38^ LBE (EE) and ml LBE (ME) in the concentration of 1250 µg/ml showed 83.61±9.98 and 72.44±5.78 % of CaSki cell inhibition after 24 h and the IC50 values of these extracts against CaSki were 220 and 490 µg /ml respectively. The IC_50_ values of water extract (WE) and ME of *Calocybe indica* in MHH-ES-1 cell line were 55.25±1.201mg/ml and 46.56±0.134mg/ml respectively while these values of MCF7 cell line were 52.12±0.15mg/ml and 47.94±0.09mg/ml respectively^39^. The IC_50_ values of EE of *Phellinus igniarius* at 48 h were 72 and 103 µg/ml for SK-Hep-1 cells and RHE cells, respectively^39^. Recently Wu *et al*.^40^ investigated that the ethanol extract of *F. pinicola* showed stronger inhibitory effect on cancer cells than water extract. Several workers reported chromatin condensation and DNA fragmentation of different cancer cells by different mushroom extracts such as EAC extract (10 g/ml) of *Cyathus striatus* treatment on HPAF-II and PL45 cells^41^, the extract of *Agaricus blazei* against leukemia NB-4 and K-562 cells^42^, and extract of *A. bisporus* on CaSki cells after 48 h treatment^38^. Treatment of COLO-205 cells with of three mushrooms (*Auricularia polytricha, Macrolepiota procera and Pleurotus ostreatus*) revealed the characteristic pattern of DNA laddering on agarose gel electrophoresis^43^. Our DNA laddering assay result is corroborated with other’s results.

Apoptosis, a genetically regulated form of cell death, plays an important role in homeostasis condition in our body^44^. Its defects can contribute to diseases like cancer, so interest surrounding the apoptosis has grown highly among oncologists. Apoptosis is suggested to be one of the major modes of action of chemotherapeutic anti-cancer drugs on malignant cells^45, 46^;there are two types of pathways in apoptosis i.e. the extrinsic pathway and the intrinsic mitochondria-mediated pathway. This induction of apoptosis was coupled with generation of reactive oxygen species (ROS), mitochondrial dysfunction, activation of caspases, and cleavage of poly (ADP-ribose) polymerase protein^47^. No mechanistic study has been conducted to investigate anti-cancer mechanism of LBE of *L. betulina* mushrooms by any previous authors. For the first time, in our study LBE of this mushroom showed the very promising anticancer effect by inducing apoptosis through ROS generation, reduction of MMP, P^53^up regulation and Bcl2, pro-caspase-3,pro-caspase 9 and PRAP down regulation. But EE of other mushrooms such as *Pleurotus ferue* on B16F10 (melanoma), BGC 823 (gastric cancer) and the immortalized human gastric epithelial mucosa cell line GES-1^48^. Comparison of the effects of extracellular, intracellular polysaccharides, and ethanol extracts of *Tremella mesenterica* revealed that only ethanol extract triggered apoptosis in A549 cells. Such effect was happened by activating a mitochondrial pathway: reduction of mitochondrial Trans membrane potential, the production of ROS (reactive oxygen species), and the activation of caspase-3 protein in ethanol extract-treated A549 cells^49^. The effects of ethanol extracts of other mushrooms on many cancer cell lines for inhibition of cell proliferation and expression of cleaved-caspase 3 and PARP, were recorded ^50, 51, 40^.

According to Xue^50^, the intra-cellular ROS generation in cancer cells by agents is an important phenomenon in cancer management. Available literatures show the anti-proliferative effects of intracellular ROS against cancer cells. As per the results of experiment conducted^52^, in ALVA-41 and PC-3 prostate cancer cells, intracellular ROS increased caspase-3 activity and cytochrome-c secreting from mitochondria trigger apoptosis. Induction of intracellular ROS by drugs e.g. cisplatin, and paclitaxel has been used to treat cancers that have high ROS status themselves ^50, 53-55^. Xue^50^ observed dramatically increased levels of extra- and intra-cellular ROS in LNCaP cells within 3 h of BBEA treatment and it triggered apoptosis. In our experiment treated cells showed higher intracellular ROS formation in contrast to untreated cells. Our results regarding ROS formations are at per with others finding but in different cell line by different mushroom extract. No scientist observed the intracellular ROS production by EE of LB in HeLa cell lines. Cell cycle analysis demonstrated that, LBE arrested cell cycle at G2/M checkpoint of HeLa cells it was also supported by our previous work where cycle was arrested at the G_2_/M checkpoint by EE of *Pluerotus florida*^56^. Similarly Jiang & Silva^57^, observed that methanolic extract of MC(mycocomplex) induced significant cell cycle arrest at G2/M phase. Moreover, cell cycle arrest at G2/M was induced by *A. blazei* in gastric epithethelial cells^58^ where as methanolic extract of *Pleurotus ostreatus* induced cell cycle arrest at G0/G1 of breast cancer (MDA-MB-231 and MCF-7) and of colon cancer cells (HCT-116 and HT-29) ^59^. Similarly Wang *et al* ^60^ found LBE of *P. ferulae* arrested cell cycle of melanoma cancer B16F10 cells at G0/G1. Interestingly, different extract from *G. lucidum* demonstrate specific effects on cell cycle progression. Thus, extracts from *G. lucidum* blocked cell cycle at G0/G1 check point in breast cancer cells^61-63^. Whereas arrest at G2/M phase was induced in prostate^62^, hepatoma^64^ and bladder cancer cells ^65^. On the other hand, Dudhgaonkar *et al*.^66^ isolated triterpenes from *G. lucidum* and observed that they triggered cell cycle arrest at G0/G1 and G2/M phase in macrophages; therefore it suggested that cell cycle arrest depends on the specific biologically active compounds as well as particular cells.

We also examined activity of the LBE as anti-metastasis by performing cologenic assay and anti-migration assay against HeLa cell line. Ganoderic acids (GA-A and GA-H) from *Ganoderma lucidum* demonstrated inhibition on colony formation of invasive breast cancer cells^67^. *Pleurotus ostreatus* (aqueous extract) exhibited a reduction in the number of colonies of COLO-205 (oral cancer) cells with 43.8 ± 3.5% in contrast with 100% proliferation on untreated cells^68^. Our Cologenic assay also showed anti colonization property as indicated by other workers in other cell line. The migration property of cancer cells is another hall mark of metastasis. In our migration scratch assay, LBE showed very good anti migration agent against HeLa cell line. It was also supported by F actin filament polymerization caused by LBE in our Laser Scanning Confocal microscopy (LSCM) study. Actin polymerization status as measured by an increase in F/G-actin ratio (increased F-actin or decreased G-actin), is a marker of cancer ^69^. Our current experiment showed that cytoplasmic F-G actin was depolymerized by LBE in HeLa cells as seen under LSC Microscopy. Lu *et al.*^65^ observed that *Ganoderma* extracts (ethanolic) triggered anti proliferation, arrested cell cycle at G2/M, and induced actin polymerization in bladder cancer cells and mentioned the impact of actin polymerization in cell migration. Since actin polymerization is a cell cycle-related event^70^, we may infer that the observed inhibition of cell proliferation and arrest in G2/M check point and inhibition of cell migration by LBE in HeLa cells may be linked with actin remodeling. However, to draw the exact relationships between actin remodeling, G 2/M arrest, and growth inhibition need more elaborated research. As we know, cytoplasmic actin filament forms podia of HeLa cells by which cells migrate and their polymerization/ shrinkage invariably inhibits podia extension and migration.

Recently autophagy has gained much attention as the potent alternative mechanism to fight cancer. Although the impact of autophagy on cancer cell death is a topic of intense debate^71, 72^, one school of oncologists with their data supports the alternate mechanism of autophagy in the death of cancer cells ^73-75^. There are emerging evidences that autophagy, a caspase-independent cell death process, plays critical roles in the generation of antineoplastic responses and mediates caspase-independent malignant cell death^76^. Other scientists studied that the deletion of Atgs and Bif-1(autophagy-regulating genes) also validated that autophagy suppressed tumor functions^77, 78^. We examined first that LBE also induced autophagy on the HeLa cell that is caspase independent cell death process. Curcumin triggered the occurrence of autophagy in SUP-B15 cells at exact 4 h and 8 h after treatment of it ^79^. Similarly, we infer that in addition to apoptosis, autophagy is another mechanism of LBE to inhibit the proliferation of HeLa cell line and tumerogenesis in mice as supported from finding of Santi *et al*^74^ who recorded that metformin prevents cell tumorigenesis through autophagy-related cell death.

Currently analysis of cancer stem cell has gained impetus in therapeutic study, as whether therapeutic strategy is able to kill all cancer stem cells (CTC) in tumor or not. Otherwise cancer/tumor will relapse after time being. So population of cancer stem cell analysis is new addition in cancer biology. We also estimated the population of stem cell of HeLa after treatment by LBE and our results showed that treatment of LBE significantly reduced CTC population after the time. A study of mice in vivo with implanted BGC-823 cancer cells revealed that the weight and volume of the tumors were reduced after 15 days’ treatment with *P. ostreatus* mycelium polysaccharide 2 (POMP2)^80^. In another study, polysaccharides were extracted from *Lepista sordida*, and when applied against laryngo carcinoma, these inhibited cell growth in vitro and caused a reduction in tumor size in vivo^81^ The compound ergosterol was isolated from *Agaricus brasiliensis*, and led to a retardation in tumor growth in sarcoma 180-bearing mice. In studies *in vivo,* ergosterol was shown to inhibit neovascularization, as well as being potentially an inhibitor of angiogenesis^82^ A marked reduction in tumor fluid volume, and changes in various other properties, were identified in Dalton’s lymphoma ascites-bearing mice following *Calocybe indica* (milky mushroom) treatment, in comparison with negative control group mice^83^ In the present study, 50 mg LBE /kg body weight of mice also led to a marked reduction in tumor volume in HeLa cell-implanted mice. None of the previous study found mechanistic clue of compounds exist in EE of mushrooms. GC-MS profile of LBE exhibited that out of 69, 14 compounds were in higher proportion with the percentage peak area > 1. Two of the 14 compounds, 9, 12-Octadecadienoic acid (Z, Z) and Ergosta-5, 8, 22-trien-3-ol, (3.beta22E) were in a significantly higher proportion with the percentage peak area 22.13 and 19.72 respectively. We started to search data base (Swiss target prediction, www.Swisstargetpridiction.ch) to find out the binding affinity cites of these two compounds. We found that Ergosta-5, 8, 22-trien-3-ol, (3. beta22E) has maximum binding affinity to Cytochrome P 450 of mitochondria, so it can attach with later and induces it for further regulation of caspase 9 apoptotic gene. Similarly, 9, 12-Octadecadienoic acid (Z, Z) has affinity to bind with fatty acid binding protein of membrane, peroxisome proliferator activated receptor, and tyrosyl DNA phosphodiesterase. The ergo sterol of Amauroderma rude, when injected in murine cancer infected mice, enhanced survival time of the mices, indicating that ergosterol might be the leading anticancer substance^84^. The inhibitors of COX-1 and COX-2 enzymes are the compounds - ergosterol, ergosta- 4,6,8^85^, 22-tetraen-3-one and 1-oleoyl-2-linoleoyl-3-palmitoylglicerol were isolated from *Grifola frondosa*^86,22^. In our LBE has much % of peak area of Ergosterol, so it conferred anti HeLa. In LBE another important fatty acid compound N-hexadecanoic acid is present and it showed significant cytotoxicity against human colorectal carcinoma cells (HCT-116)^87^. Harada *et al*.^87,88^ reported that N-hexadecanoicacid blocked proliferation of human fibroblast cells by inhibiting DNA topoisomerase-I. Lai *et al*.^89^ reported that it binds to microtubule associated protein ‘tau’, and inhibits cell division. Other workers demonstrated that both oleic and α –linolenic acid showed a proliferation inhibition effect on prostate carcinoma cells^90, 91^. More over phenolic compounds such as Phenol, 2,4-bis(1,1-dimethylethyl) (1.50) and Phenol, 2,4,6-tris (21icycle[2.2.1]h ept-2-yl) are also found in LBE. Role of phenolic compounds as anticancer is now well established. A recent study showed that 2, 4-bis (1,1-dimethylethyl)-(1) is cytotoxicity against Hela cell^92^. Mechanistic studies have shown that phenolic compounds can produce reactive oxygen species (ROS) under cell culture conditions and induce apoptosis in cancer cells^93-98^. It has been reported that polyphenolic compounds have inhibitory effects on mutagenesis and carcinogenesis in humans, when up to 1.0 gm is ingested daily from a diet rich in fruits and vegetables^99^. Only four phenolic acids, however, were analyzed in *Agaricus* (white &brown) and shiitake mushrooms^100^. In our experiment LBE showed potent anti proliferative property with IC_50_ at 492.52±2.6 µg/ml, 612.22 ± 4.2 µg/ml, and 1210.30 ± 6.4 µg/ml for 24 h against HeLa, CaSki and SiHa cell line respectively. Liu *et al*. ^101^ isolated eight compounds including 5 sterols from *L. betulina,* while Fakoya & Oloketuyi^9^, reported phenolic and steroids compounds are present in EE of this mushroom. Lupeol, one terpenoid, is also present in LBE, library search exhibited that it facilitated significant inhibition of growth of breast cancer cell line^102^, prostate^103^, colorectal^104^, and gastric cancer^105^. GCMS profile of LBE also showed the presence of 4’-Hydroxy-6-methoxyaurone and this compound inhibits cancer cells to acquire drug resistant and also cytotoxicity against mouse RAW264.7 cell^106^. Recently multidrug resistance (MDR) of tumor cells to cytotoxic drugs is a major problem in cancer chemotherapy^106^. It has been well established that the mechanism by which tumor cells acquire multidrug resistance (MDR) is the over expression of aPgp (P-glycoprotein: ATP dependent membrane glycoprotein) that binds a drug and drags out it from the cell^107-109^. The driving force of drug efflux is Pgp-mediated ATP hydrolysis, the liberated energy of which is used for the drug transport^110-112^. 4-Hydroxy-6-methoxyaurone has high binding affinity to the nucleotide-binding domain of Pgp^113^ and by binding with Pgp it may inhibit the cancer cells to become drug resistant.^112^

In conclusion, LBE (LBE of *L.betulina*) is a mixture of potential compounds for inhibition of proliferation of HeLa, CaSki and SiHa cell lines. By apoptosis as shown by ROS generation, MMP reduction, regulating pro-caspase 3 & 9, and cancer guard gene p53, it was found that LBE is potent anti-HeLa cell line. Moreover it induced caspase independent autophagy of HeLa cells and prevents tumerogenesis. It blocked the cell division by arresting G2/M checkpoint. Anti-metastasis property of it was also revealed from anti-migration and anti-cologenic capacity. Its anti-migration property was also validated from F-G actin polymerization capacity in HeLa cells. Efficacy to reduce cancer stem cell population is one another important anti-HeLa property of this mushroom. Antitumor effects of LBE treatment in HeLa-implanted mice model, in our experiment also validated that LBE is one important anti cervical cancer activity. GC-MS profile also exhibited a lot of anti-cancerous fatty acids, phenol and terpenoid. One interesting compound i.e. 4’-Hydroxy-6-methoxyaurone which inhibits the cancer cells to become drug resistant. Taken together, the anticancer activity of ethanol extract of *L. betulina* may be the result of the synergistic effects of various compounds, which suggests that this mushroom is excellent source of anticancer agents in the treatment of cervical cancer.

## Acknowledgements

Authors acknowledge National Test House, Kolkata for providing GC-MS facility and principle of Ramakrishna Mission Vivekananda Centenary College (Autonomous) for providing laboratory facilities. Authors are very grateful to Professor Ananda Mohan Chakrabarty for giving needful suggestion for designing of the research.

## Funding information

Authors acknowledge the financial support from the Department of Science & Biotechnology, GoWB, W.B., India. (No.840 (sanc)/ST/P/S&T/1G-11/2015). The funders had no role in study design, data collection and analysis, decision to publish, or preparation of the manuscript.

## Author’s contribution

S.K.G. has done research design and written manuscript. T.S. has done experimental works, data collection and data analysis.

## Data availability statement

All relevant data are within the manuscript

## Competing financial and nonfinancial interests

Authors declare no competing financial and nonfinancial interest.

